# JAG1 intracellular domain acts as a transcriptional cofactor that forms an oncogenic transcriptional complex with DDX17/SMAD3/TGIF2

**DOI:** 10.1101/2021.03.31.437839

**Authors:** Eun-Jung Kim, Jung Yun Kim, Sung-Ok Kim, Seok Won Ham, Sang-Hun Choi, Nayoung Hong, Min Gi Park, Junseok Jang, Sunyoung Seo, Kanghun Lee, Hyeon Ju Jeong, Sung Jin Kim, Sohee Jeong, Kyungim Min, Sung-Chan Kim, Xiong Jin, Se Hoon Kim, Sung-Hak Kim, Hyunggee Kim

**Affiliations:** Department of Biotechnology, College of Life Sciences and Biotechnology, Korea University, Seoul 02841, Republic of Korea; Institute of Animal Molecular Biotechnology, Korea University, Seoul 02841, Republic of Korea; Department of Biochemistry, College of Medicine, Hallym University, Chuncheon 24252, Republic of Korea; School of Pharmacy, Henan University, Kaifeng, Henan 475004, China; Department of Pathology, Severance Hospital, Yonsei University College of Medicine, Seoul 03722, Republic of Korea; Department of Animal Science, College of Agriculture and Life Sciences, Chonnam National University, Gwangju 61186, Republic of Korea; Gwangju Center, Korea Basic Science Institute, Gwangju 61186, Republic of Korea

**Author notes:** These authors contributed equally to this work (Eun-Jung Kim, Jung Yun Kim).

## Abstract

Jagged1 (JAG1) is a Notch ligand that contact-dependently activates Notch receptors and regulates cancer progression. The JAG1 intracellular domain (JICD1) is generated from JAG1, such as the formation of NOTCH1 intracellular domain (NICD1), however, the role of JICD1 in tumorigenicity has not been comprehensively elucidated. Herein, we revealed that JICD1 induced astrocytes to acquire several cancer stem cell properties, including tumor formation, invasiveness, stemness, and resistance to chemotherapy and radiotherapy. The transcriptome, ChIP-sequencing, and proteomic analyses revealed that JICD1 increased SOX2 expression by forming a transcriptional complex with DDX17, SMAD3, and TGIF2. Furthermore, JICD1-driven tumorigenicity was directly regulated by SOX2. Therefore, our results demonstrated that, like NICD1, JICD1 acts as a transcriptional cofactor in the formation of the DDX17/SMAD3/TGIF2 transcriptional complex, leading to oncogenic transformation.

## Introduction

Glioblastoma (GBM) is reportedly the most malignant brain tumor, classified as World Health Organization (WHO) grade IV, presenting characteristics such as rapid cell proliferation, diffusive margins, aberrant vascularization, and heterogeneous populations. Despite the use of post-operative chemotherapy and radiotherapy, the median survival time for patients with GBM is less than 14 months^1^. One of the obstacles in the treatment of GBM is GBM stem cells (GSCs), which is a small population of tumors. GSCs are known to demonstrate high tumorigenicity, self-renewal, and resistance to anticancer therapy^2, 3, 4^. Accordingly, strategies for eliminating GSCs are of considerable interest in basic and clinical research.

The Notch signal regulates cell fate determination during embryonic development and in adult tissue homeostasis and is associated with tumor formation and progression^5, 6^. In particular, Notch signaling plays an important role in the acquisition and maintenance of cancer stem cell characteristics^7, 8^, leading to intensive research in recent decades.

The Notch signaling pathway is a contact-dependent signaling pathway and is evolutionarily highly conserved. In vertebrates, Notch signaling components consist of four receptors (NOTCH1 to 4) and five ligands [delta-like 1 (DLL1), DLL3, DLL4, and jagged-1 (JAG1), JAG2]^9^. The Notch receptors are type I transmembrane glycoproteins that are transported to the cell membrane after s1 cleavage by a furin-like convertase in the Golgi apparatus. In the canonical ligand-dependent pathway, Notch receptors undergo structural transformation upon ligand engagement, with s2 cleavage in the extracellular domain mediated by metalloprotease ADAM10/17 (A disintegrin and metalloproteinase domain-containing protein 10/17). In turn, the s2 cleavage triggers s3 cleavage by γ-secretase within the transmembrane domain of receptors. The consecutive proteolytic cleavage generates the Notch receptor intracellular domain (NICD), which translocates to the nucleus^10^. NICD cannot directly bind to DNA and interacts with the DNA-binding protein CSL (CBF1/RBPJ, Suppressor of hairless, Lag-1)^11, 12^. CSL can recruit both the transcription activator and repressor. In the absence of NICD, CSL interacts with the transcription repression complex. The NICD/CSL complex activates the expression of canonical Notch target genes by recruiting co-activators, including Mastermind-like protein family, p300, histone acetyltransferases, and chromatin remodeling factors^13, 14^.

JAG1, a Notch ligand, has been introduced as a target of anticancer therapy owing to its crucial role as an oncogene in a wide array of solid tumors, as well as in the formation of cancer stem cells^15, 16^. However, to date, the role of JAG1 has been limited to the activation of Notch receptors. Interestingly, although JAG1 expression was increased in accordance with the increasing malignancy in patients with glioma, the expression of cleaved NOTCH1, which is activated by JAG1, was reportedly decreased in patients with grade IV GBM patients^16, 17^. This indicates that JAG1 possesses a secondary signaling mechanism, independent of the signal that activates the Notch receptors, to promote malignancy.

Furthermore, several studies have revealed that Notch ligands release the intracellular domain (ICD) with identical processing as NICD formation^18, 19^. The extracellular domain of full-length JAG1 is cleaved by ADAM10/17, resulting in soluble JAG1 leaving the cells, with the remaining C-terminal fragment (CTF) sequentially cleaved by γ-secretase. Subsequently, this cleavage generates the JAG1 intracellular domain (JICD1), and JICD1 migrates into the nucleus, like NICD. However, it remains unclear what kind of transcription factor is associated with JICD1 and how JICD1 regulates target gene expression.

In this study, we focused on two aspects of the JICD1 function: oncogenic and transcriptional properties. Our data indicated that JICD1 functions as a transcriptional cofactor in cooperation with DDX17, SMAD3, and TGIF2, and promotes tumorigenesis by acquiring GSC properties through SOX2 expression.

## Results

### JAG1 generates JICD1 that is localized in the nucleus of glioblastoma cells

To determine the expression level of JAG1 in brain tumors, we analyzed the expression of Notch receptors and ligands using the Repository for Molecular Brain Neoplasia Data (REMBRANDT) database^20^. The expression levels of Notch receptors and ligands differed depending on the tumor type, WHO grade, degree of DNA methylation, and GBM subtype (Fig. 1a). On comparing the expression levels of JAG1 and NOTCH1, our findings revealed that JAG1 expression increased with increasing WHO grade, and NOTCH1 expression was lower in WHO grade IV than that in grades II and III (Supplementary Fig. 1a). GBM is composed of heterogeneous populations and is classified into four subtypes: proneural, neural, mesenchymal, and classical^21^. The characteristics of the neural subtype are similar to normal brain tissues, but the mesenchymal and classical subtypes are more aggressive than that is the proneural subtype^22^. The proneural subtype is enriched in Notch-related factors^23^. Moreover, NOTCH1 is highly expressed in the proneural subtype, whereas its expression progressively decreases in mesenchymal subtypes. In contrast, JAG1 is highly expressed in the classical, mesenchymal, and proneural subtypes, in the stated order, but its expression is lower in neural subtypes than that in other subtypes (Supplementary Fig. 1b). Thus, JAG1 and NOTCH1 were differentially expressed depending on tumor malignancy and subtype. Furthermore, although NOTCH1 expression does not affect patient prognosis, the high JAG1-expressing group presented poor survival rates among patients with glioma (Fig. 1b).

**Fig. 1:**
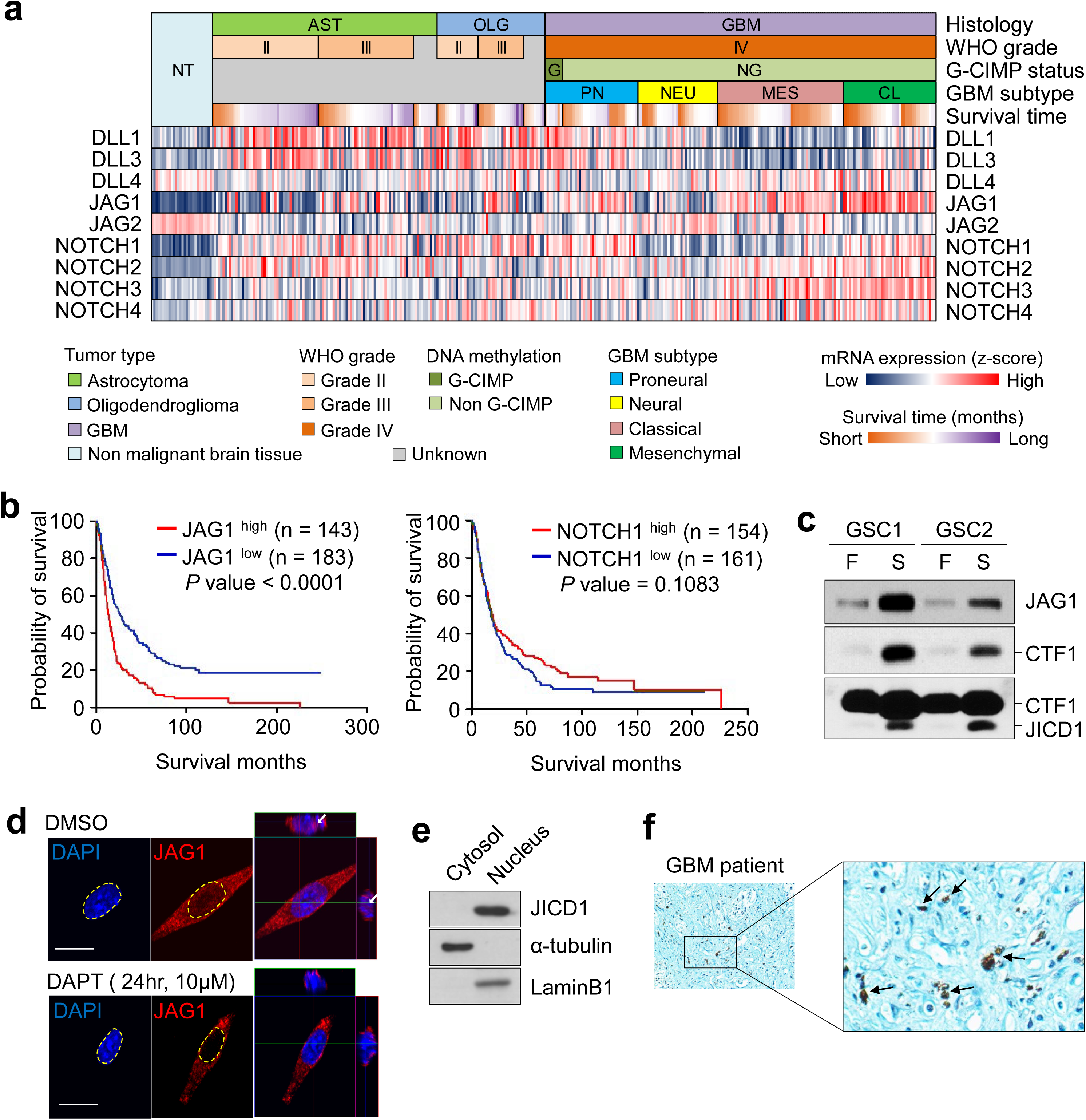
JAG1 produces JICD1 that translocates into the nucleus in GBM stem cells (GSCs). **a** Heatmap showing the expression of Notch ligands and receptors based on histology, WHO grade, G-CIMP status, GBM subtype, and survival time using the REMBRANDT database. **b** Survival time of patients with glioma according to expressions of JAG1 (left) and NOTCH1 (right). The patients were divided into two groups (high and low) based on their expression (mean ± S.E.M). *p*-value with a log-rank (Mantel-Cox) test. **c** Expression of JAG1, CTF1, and JICD1 in GSCs grown in the stem cell (S: DMEM/F12 with B27, EGF, and bFGF) and differentiation (F: DMEM with FBS) culture conditions. **d** Expression of JICD1 in HEK293T cell after γ-secretase inhibitor (DAPT) treatment. The red fluorescent signal indicates JAG1, blue-DAPI was used to stain nuclei. Scale bar = 20 µ m. Arrows refer to nuclear staining of JAG1 (JICD1). **e** Expression of JICD1 in the nuclear and cytoplasmic fractions of HA-JICD1-overexpressing HEK293T cells. LaminB1 and α-tubulin were used as nuclear and cytoplasmic fraction markers, respectively. **f** Expression of JAG1 in human GBM specimen. Arrows indicate nuclear staining of JAG1 (JICD1). See also Supplementary Fig. 1.

To further elucidate the role of JAG1 in glioma biology, we examined the expression of JAG1 in patient-derived GSCs grown under stem cell or differentiation culture conditions. JAG1 expression was higher in GSCs grown under stem cell culture conditions (Fig. 1c). Interestingly, expression of both CTF and JICD1 was also relatively higher in GSCs grown under stem cell culture conditions. Several studies have demonstrated the oncogenic roles of JAG1 in various human malignancies^24, 25, 26^. However, unlike NICD1 produced by the Notch receptor, the oncogenic role of JICD1 in tumor initiation and progression has not been comprehensively elucidated. Nuclear localization of endogenous JICD1 was observed in single cells and was inhibited by treatment with DAPT, a γ-secretase inhibitor (Fig. 1d). Cell fractionation assays demonstrated that endogenous JICD1 translocated into the nucleus (Fig. 1e). These results suggested that JICD1 is produced from JAG1 by γ-secretase without cell-cell contact. In human GBM specimens, JICD1 was predominantly localized in the nucleus (Fig. 1f). Collectively, these results suggested that, in contrast to the importance of the Notch receptor-signaling pathway in tumorigenesis and malignancy, JAG1 and JICD1 may be associated with brain tumor malignancies.

### JICD1 plays an essential role in acquiring GSC features

Notch signaling is important for tumor formation and maintenance of cancer stem cells. However, ectopic overexpression of NICD1 in Ink4a/Arf^-/-^ astrocytes (derived from mice deficient in the CDKN2A gene) promote dedifferentiation and neurosphere formation *in vitro* but are non-tumorigenic *in vivo*^27^. To compare the oncogenic function of JICD1 with that of NICD1, we established JICD1 (HA-tagging JICD1)- and NICD1-overexpressing Ink4a/Arf^-/-^ astrocytes (Fig. 2a). Cell growth was increased by NICD1, but not by JICD1 (Fig. 2b); the sphere-forming ability was equally improved in both cell types (Fig. 2c). Interestingly, intracranial injection of control, JICD1-, and NICD1-overexpressing Ink4a/Arf^-/-^ astrocytes revealed that only JICD1-overexpressing cells formed tumors (Fig. 2d). JICD1- overexpressing brain tumors invaded surrounding normal tissues, and JICD1-overexpressing cells were also invasive in the Matrigel invasion assay (Fig. 2e, f). Furthermore, JICD1 induced resistance to the anticancer agents, temozolomide (TMZ) and carmustine (BCNU), whereas resistance to radiation was induced by both JICD1 and NICD1 (Fig. 2g and Supplementary Fig. 2). Collectively, these results suggested that JICD1 plays a significant role in tumorigenesis, invasiveness, and resistance to anticancer therapy.

**Fig. 2:**
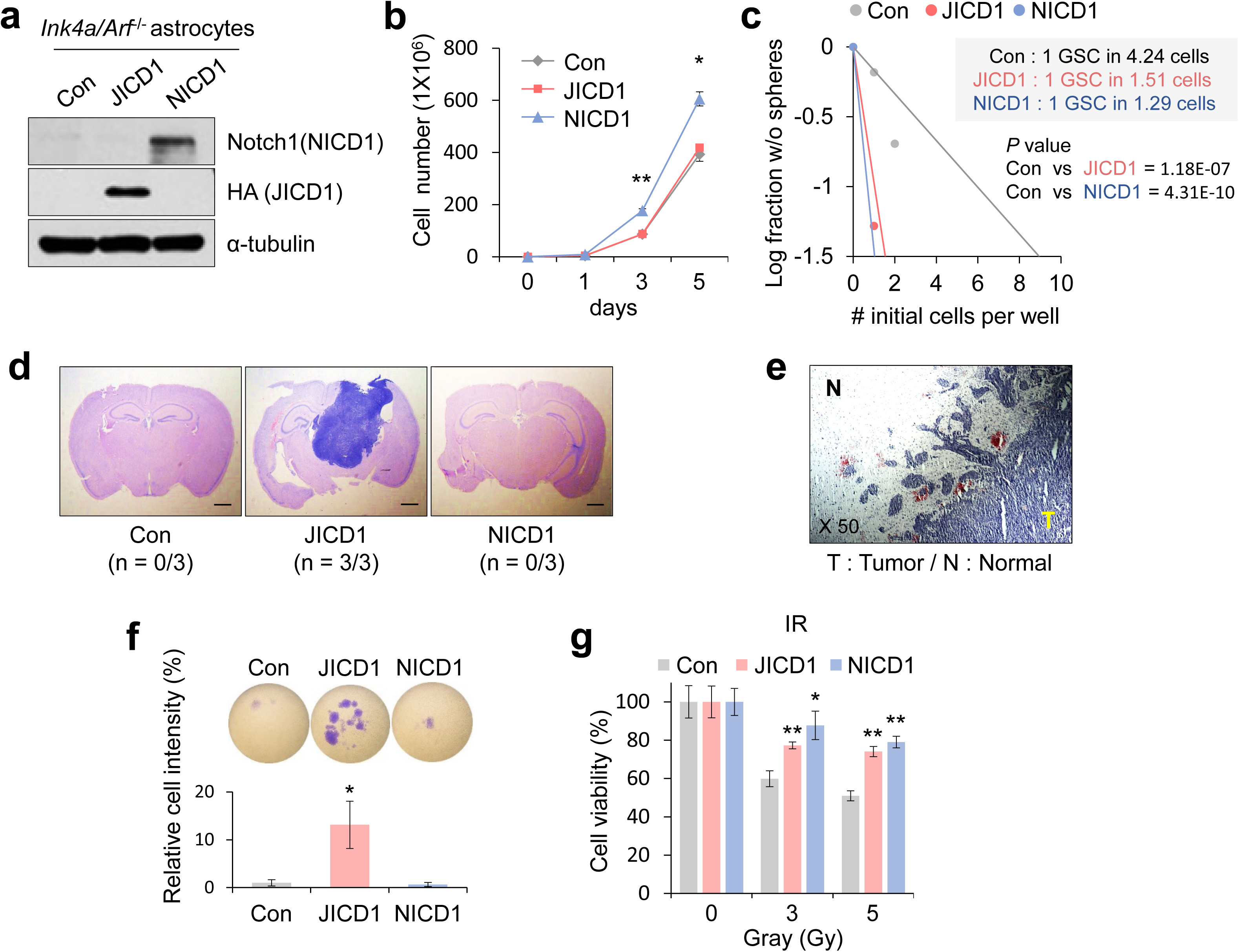
JICD1 promotes acquisition of GSC properties. **a** Ectopic expression of HA-JICD1 and NICD1 in murine Ink4a/Arf^-/-^ astrocytes. **b** Cell growth of control, HA-JICD1- and NICD1-overexpressing Ink4a/Arf^-/-^ astrocytes. \**p* < 0.05, \*\**p* < 0.01. **c** Sphere forming ability of control, HA-JICD1- and NICD1-overexpressing Ink4a/Arf^-/-^ astrocytes determined by an *in vitro* limiting dilution assay. **d** Representative H&E staining showing mouse brain sections after intracranial injection of control, HA-JICD1- and NICD1-overexpressing Ink4a/Arf^-/-^ astrocytes. Scale bar = 500 µ m. **e** The marginal region of mouse brain tumor derived from HA-JICD1- overexpressing Ink4a/Arf^-/-^ astrocytes. Magnification 50×. (T: Tumor, N: Normal). **f** *In vitro* invasion ability of control, HA-JICD1- and NICD1-overexpressing Ink4a/Arf^-/-^ astrocytes for 48h. \**p* < 0.05. **g** Irradiation resistance of control, HA-JICD1- and NICD1-overexpressing Ink4a/Arf^-/-^ astrocytes over 48h. \**p* < 0.05, \*\**p* < 0.01. Data were obtained from three independent experiments (mean ± S.E.M). See also Supplementary Fig. 2.

### JICD1 exerts oncogenic functions by regulating SOX2 expression

To clarify how JICD1 regulates GSC features and oncogenic effects, we performed RNA-sequencing (RNA-Seq) in control, JICD1-, and NICD1-overexpressing Ink4a/Arf^-/-^ astrocytes. Compared with that by the control, JICD1- and NICD1-overexpressing cells upregulated 1024 and 1291 genes, respectively. Of these, 488 genes were commonly upregulated, and 536 genes were specifically upregulated in JICD1-overexpressing cells (Fig. 3a). The characteristics of JICD1- and NICD1-overexpressing cells could be determined using these differentially expressed genes (DEGs). To determine target genes that confer the JICD1 characteristics, first, the gene ontology (GO) of JICD1-upregulated DEGs (n=536) was analyzed using the Database for Annotation, Visualization and Integrated Discovery (DAVID) bioinformatics resource^28^. The biological process (BP) of JICD1-upregulated DEGs showed enrichment for biosynthetic process, chromatin organization, cell motility, and development (Supplementary Fig. 3a). The enrichment of these BPs was consistent with the observed JICD1 characteristics (Fig. 1 and 2). The cellular component (CC) of JICD1-upregulated DEGs is found to be associated with chromatin, highlighting the role of JICD1 in the nucleus (Supplementary Fig. 3b). Patients with glioma presenting high levels of JICD1-upregulated DEGs had shorter survival times than those with lower levels of these DEGs, indicating the clinical relevance of JICD1-upregulated DEGs (Fig. 3b). Among the significant GOs, we selected genes that positively correlated with JAG1 expression, were expressed highly in tumors when compared that in with non-tumors and presented a poor prognosis for patient survival from the REMBRANDT database (Fig. 3c and Supplementary Fig. 3c-e). We identified *Pou3f2* (POU Class 3 Homeobox 2), *Sox2* (Sex determining region Y-box 2), and *Mreg* (Melanoregulin) genes as putative regulators of JICD1 characteristics, and confirmed that their expression increased in JICD1- overexpressing cells (Fig. 3d). Additionally, 569 JICD1-downregulated DEGs were identified, and BPs of these DEGs showed enrichment for cell adhesion (Supplementary Fig. 4a, b). The analysis resulted in the identification of genes *Omd*, *Pcdh19*, and *Wnt7b*, which that inversely correlated with glioma grade, patient prognosis and JAG1 expression (Supplementary Fig. 4c-f). Furthermore, BP analysis was performed on DEGs upregulated or downregulated by NICD1, and on DEGs upregulated by both JICD1 and NICD1 (Supplementary Fig. 5). BP of DEGs, which are upregulated or downregulated by NICD1, involved biosynthetic process, development, immune response and cell adhesion (Supplementary Fig. 3a, 4b, and 5a-b). However, this is inconsistent with the characteristics of NICD1-overexpressing cells (Fig. 2).

**Fig. 3:**
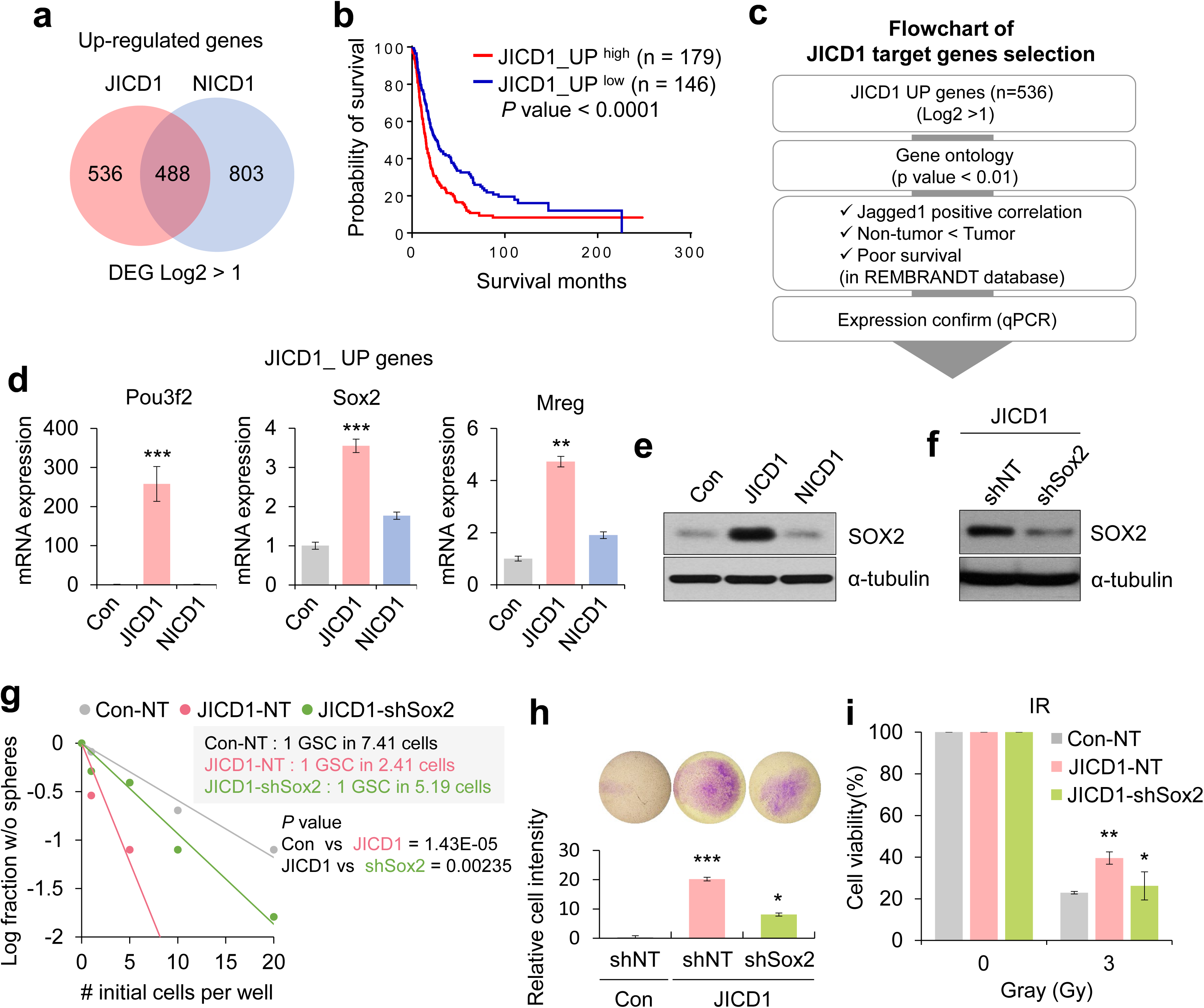
JICD1 modulates the characteristics of GSCs by regulating SOX2 expression. **a** Venn diagram showing upregulated DEGs in HA-JICD1- and NICD1-overexpressing Ink4a/Arf^-/-^ astrocytes. Fold-change > 2. **b** Correlation between glioma patient survival and JICD1-upregulated DEG enrichment score in the REMBRANDT database. The patients were divided into two groups (high and low) based on their JICD1-upregulated DEG enrichment score (mean ± S.E.M). **c** A flowchart showing the process of selecting JICD1 target genes. **d** The relative mRNA expression of JICD1 target genes (*Pou3f2*, *Sox2*, and *Mreg*) in control, HA-JICD1-, and NICD1-overexpressing Ink4a/Arf^-/-^ astrocytes. \*\**p* < 0.01, \*\*\**p* < 0.001. **e** Expression of SOX2 protein in control, HA-JICD1-, and NICD1- overexpressing Ink4a/Arf^-/-^ astrocytes. α-Tubulin was used as a loading control. **f** Expression of SOX2 protein in JICD1-overexpressing Ink4a/Arf^-/-^ astrocytes after shSox2 infection. NT indicates infection with the non-target, control shRNA. **g** Sphere forming ability of control-shNT, JICD1-shNT, and JICD1-shSox2 Ink4a/Arf^-/-^ astrocytes determined by an *in vitro* limiting dilution assay. **h** *In vitro* invasion ability of control-shNT, JICD1-shNT, and JICD1-shSox2 Ink4a/Arf^-/-^ astrocytes for 48 h. \**p* < 0.05, \*\*\**p* < 0.001. **i** Irradiation resistance of control-shNT, JICD1-shNT, and JICD1-shSox2 Ink4a/Arf^-/-^ astrocytes over 48 h. \**p* < 0.05, \*\**p* < 0.01. Data were obtained from three independent experiments (mean ± S.E.M). See also Supplementary Fig. 3-5.

Among JICD1-upregulated DEGs, SOX2 plays a pivotal role in the maintenance of embryonic and neural stem cells and is associated with cancer progression in numerous solid tumors^29, 30, 31^. To confirm the oncogenic function of SOX2, we established SOX2-overexpressing cells (Supplementary Fig. 6a). As expected, SOX2 induced tumor characteristics such as sphere formation, invasion, and resistance to chemotherapy (Supplementary Fig. 6b-e). Expression of SOX2 protein was increased in JICD1- overexpressing cells (Fig. 3e). To clarify whether the GSC properties acquired by JICD1 can be attributed to increased SOX2 expression, we established SOX2-depleted cells in JICD1-overexpressing cells using a short hairpin RNA targeting *Sox2*^31^ (Fig. 3f). The sphere-forming ability was significantly decreased by SOX2 depletion in JICD1-overexpressing cells (Fig. 3g). Furthermore, the invasion ability and radiation resistance decreased (Fig. 3h, i). Collectively, these results indicated that SOX2 plays a crucial role in regulating the oncogenic function of JICD1.

### JICD1 regulates SOX2 transcription in cooperation with DDX17

In the canonical Notch signaling pathway, NICD interacts with the CSL transcription factor to regulate Notch target genes^11^. To determine whether JICD1 is also capable of interacting with CSL, a CSL- luciferase reporter assay was performed. In contrast to NICD1, JICD1 did not activate the CSL promoter activity (Supplementary Fig. 2c). Next, we performed cell fractionation and mass spectrometry to identify JICD1 binding factors (Supplementary Fig. 7a). This study showed that JICD1 interacts with 21 proteins in the nucleus and 26 proteins in the cytoplasm (Fig. 4a and Supplementary Fig. 7b). Most JICD1 binding proteins positively correlated with JAG1 in the REMBRANDT database. DDX17 (DEAD-Box Helicase 17), HSPA5 (Heat Shock Protein Family A Member 5), and PDIA6 (Protein Disulfide Isomerase Family A Member 6) were bound with JICD1 in both the nucleus and cytoplasm. Of the 21 JICD1 binding proteins in the nucleus, only DDX17 is known to be involved in transcriptional regulation.

**Fig. 4:**
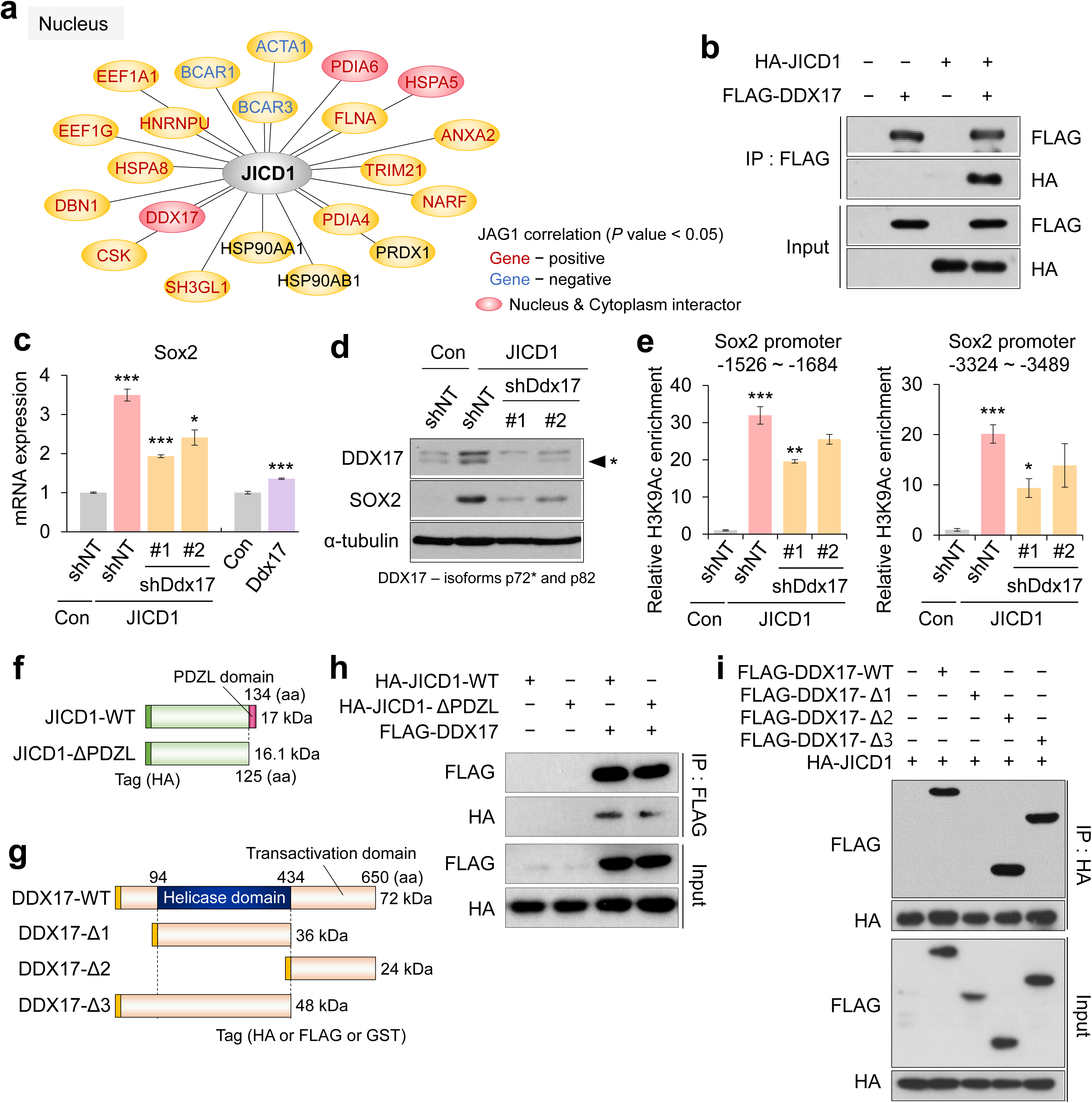
JICD1 interacts with DDX17 and activates SOX2 transcription. **a** A schematic diagram that presents JICD1 binding proteins in the nucleus identified by affinity purification and tandem mass spectrometry. Font color indicates proteins that are correlated with JAG1 in the REMBRANDT database (red: positive, blue: negative, and black: no correlation). The red circle indicates proteins that bind to JICD1 in both the nucleus and cytoplasm. **b** The association between HA-JICD1 and FLAG-DDX17 in HEK293T cells. **c** Expression of SOX2 mRNA in control-shNT, HA- JICD1-shNT, two HA-JICD1-shDdx17, control, and DDX17-overexpressing Ink4a/Arf^-/-^ astrocytes. \**p* < 0.05, \*\*\**p* < 0.001. **d** Expression of SOX2 and DDX17 proteins in control-shNT, HA-JICD1-shNT, and two HA-JICD1-shDdx17 Ink4a/Arf^-/-^ astrocytes. **e** ChIP-PCR analysis of the H3K9 acetylation status within two regions of Sox2 promoter in control-shNT, HA-JICD1-shNT, and two HA-JICD1- shDdx17 Ink4a/Arf^-/-^ astrocytes. **f**, **g** Schematic diagrams that represent JICD1 (F) and DDX17 (G) mutants. (WT: Wild-type, △: Mutant, PDZL: PSD-95/Dlg/ZO-1 ligand). **h** The association between HA-JICD1 mutants and FLAG-DDX17 in HEK293T cells. **i** The association between HA-JICD1 and FLAG-DDX17 mutants in HEK293T cells. See also Supplementary Fig.7, 8.

DDX17 unwinds RNA through ATP binding and hydrolysis and regulates RNA metabolism, including pre-RNA splicing, alternative splicing, and miRNA processing^32^. Furthermore, DDX17 controls transcription by recruiting RNA polymerase II and histone modification factors^33^. The co-transfection of JICD1 and DDX17 into HEK293T cells revealed co-localization and interaction (Fig. 4b and Supplementary Fig. 8a). As JICD1 regulates SOX2 expression and induces oncogenic function, we next evaluated whether DDX17 regulates SOX2 expression. The mRNA and protein levels of SOX2 dramatically decreased in DDX17-depleted JICD1 cells (JICD1-shDdx17 cells) when compared with the JICD1-shNT cells (non-targeting control cells) (Fig. 4c, d). Then, we investigated the molecular mechanism through which DDX17 regulates mRNA and protein expression of SOX2. Quantitative chromatin immunoprecipitation (ChIP)-PCR analysis was performed to determine histone H3 lysine 9 (H3K9) acetylation, a known transcription activation marker^34^. JICD1 enhanced H3K9 acetylation of the *Sox2* promoter, however, depletion of DDX17 was attenuated (Fig. 4e). Interestingly, SOX2 expression was not markedly altered by DDX17 overexpression alone (Fig. 4c), indicating that SOX2 transcription can be activated by the JICD1/DDX17 complex. To investigate the regions of JICD and DDX17 proteins responsible for interaction, several deletion mutants of JICD1 and DDX17 were generated (Fig. 4f, g). Deletion of the PDZL domain of JICD1 did not affect on JICD1/DDX17 binding (Fig. 4h). Deletion of the transactivation domain of DDX17 completely disrupted the interaction between JICD1 and DDX17 (Fig. 4i). These data revealed that among various JICD1 binding proteins in the nucleus and cytoplasm, JICD1 interacts with the transactivation domain of DDX17 to regulate SOX2 expression.

### JICD1/DDX17 complex binds to *Sox2* promoter via transcription factor SMAD3

DDX17 interacts with transcription factors without DNA binding and acts as an adaptor to recruit various transcriptional regulatory factors^35^. To identify transcription factors associated with the JICD1/DDX17 complex, transcription factor binding (TFB) DNA motifs of chromatin that interact with JICD1 were defined by ChIP-sequencing (ChIP-seq) analysis. Furthermore, the activated transcription-associated DNA motifs were detected by H3K9 acetylation. We identified 46 TFB DNA motifs interacting with JICD1 and 78 motifs acetylated at H3K9 in JICD1-overexpressing cells (Fig. 5a).

**Fig. 5:**
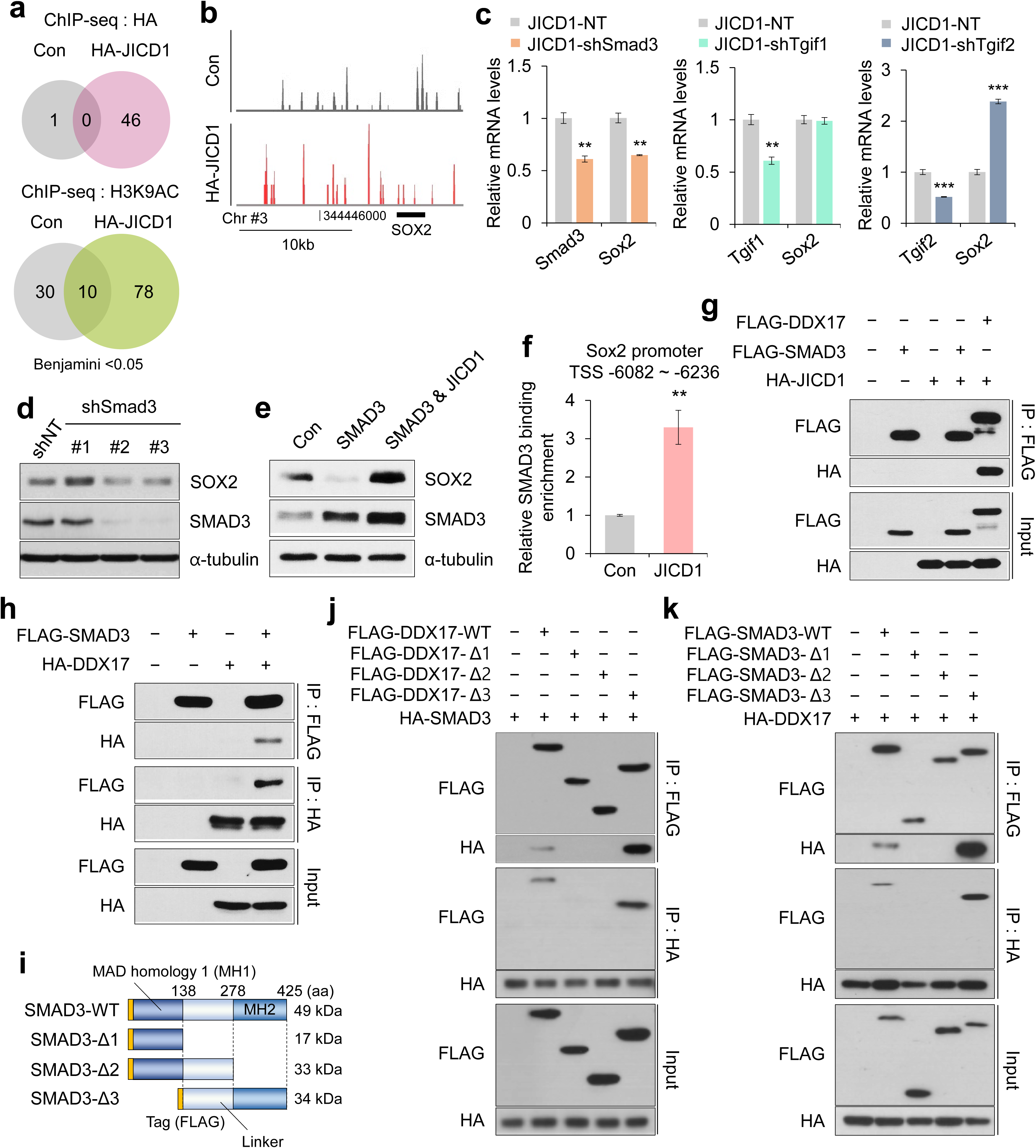
JICD1/DDX17 complex binds to DNA according to association with SMAD3. **a** Venn diagrams showing transcription factor binding DNA motifs that specifically interact with JICD1 (top) and H3K9-acetylated transcription factor binding DNA motifs (bottom) in the control and HA- JICD1-overexpressing Ink4a/Arf^-/-^ astrocytes from ChIP-seq analysis. **b** Binding intensity of anti-HA (JICD1) in the control versus HA-JICD1-overexpressing Ink4a/Arf^-/-^ astrocytes at the genomic locus of Sox2. **c** Expression of Sox2 mRNA in HA-JICD1-shNT, HA-JICD1-shSmad3, HA-JICD1-shTgif1, and HA-JICD1-shTgif2 Ink4a/Arf^-/-^ astrocytes. \*\**p* < 0.01, \*\*\**p* < 0.001. **d** Expression of SOX2 protein in Ink4a/Arf^-/-^ astrocytes after shSmad3 infection. NT indicates infection with the non-target, control shRNA. **e** Expression of Sox2 mRNA in control, FLAG-SMAD3-, FLAG-SMAD3-HA-JICD- overexpressing Ink4a/Arf^-/-^ astrocytes. **f** ChIP-PCR analysis of the SMAD3 engagement to its binding motif at the Sox2 promoter in control, HA-JICD1 Ink4a/Arf^-/-^ astrocytes. \*\**p* < 0.01. **g** Co-IP analysis between FLAG-SMAD3 and HA-JICD1. The association of FLAG-DDX17 with HA-JICD1 was used as a positive control. **h** The association between HA-DDX17 and FLAG-SMAD3 in HEK293T cells. **i** Schematic diagrams showing SMAD3 mutants. (WT: Wild-type, △: Mutant, MH: MAD homology). **j** The association between HA-SMAD3 and FLAG-DDX17 mutants in HEK293T cells. **k** The association between HA-DDX17 and FLAG-SMAD3 mutants in HEK293T cells. See also Supplementary Fig. 8, 9.

JICD1 associated TFB DNA motifs were enriched near the transcription start site (TSS) in the same manner as activated transcription-associated DNA motifs (Supplementary Fig. 9a). Indeed, JICD1 was predominantly associated with the *Sox2* promoter when compared with that in the control (Fig. 5b). Transcription factors that bind these motifs associated with JICD1 in the *Sox2* promoter were SMAD3 (SMAD Family Member 3), TGIF1 (TGFB-induced factor homeobox 1), and TGIF2 (TGFB-induced factor homeobox 2) (Supplementary Fig. 9b). To investigate whether these transcription factors regulate SOX2 expression, we depleted the expression of each transcription factor in JICD1-overexpressing cells using shRNAs (Fig. 5c). Depletion of SMAD3 decreased the mRNA and protein expression of SOX2 (Fig. 5c, d). Additionally, depletion of TGIF2 increased the expression of *Sox2* (Fig. 5c). In the presence of JICD1, SOX2 expression was increased dependent on SMAD3 (Fig. 5e and Supplementary Fig. 9c). Furthermore, SMAD3 enabled binding more predominantly to the SMAD3 binding motif on the *Sox2* promoter in JICD1-overexpressing cells than that in the control cells (Fig. 5f). However, SMAD3 reduced mRNA and protein expression of SOX2, independent of JICD1 (Fig. 5e and Supplementary Fig. 9c). Based on these observations, we concluded that the transcription of SOX2 is regulated via SMAD3 in the presence of JICD1.

SMAD3 is a receptor-regulated SMAD and transcription modulator in the transforming growth factor-beta (TGF-β) signaling pathway^36, 37^. There are two conserved domains in SMAD3: the MAD homology 1 (MH1) domain binds DNA and the MH2 domain recruits transcriptional machinery such as CBP, p300, c-Ski, and SnoN. We confirmed that SMAD3 interacted with DDX17, but not JICD1 (Fig. 5g, h) and co-localized with DDX17 in the nucleus (Supplementary Fig. 8b). To map the region interacting with DDX17, deletion mutants of SMAD3 were generated (Fig. 5i). SMAD3 was bound to the N-term of DDX17, but not to the core helicase and transactivation domains (Fig. 5j). Additionally, DDX17 bind to the MH2 domain of SMAD3, which is a well-known interaction region with other transcriptional machinery (Fig. 5k). These data revealed that SMAD3 is associated with the JICD1/DDX17 complex and that the MH2 domain of SMAD3 is crucial for mediating the interaction with the N-term of DDX17.

### SMAD3 acts as a molecular hub for interaction with the JICD1/DDX17/SMAD3/TGIF2 complex

We identified 46 TFB DNA motifs interacting with JICD1 and screened these motifs in the promoter of JICD1-upregulated DEGs (Supplementary Fig. 9d). JICD1-upregulated DEG promoters have several TFB DNA motifs, in the order, TGIF1, TGIF2, SOX3, SMAD3, and SMAD4. TGIF1 and TGIF2 are transcription factors that mostly bind to the promoter of JICD1-upregulated DEGs. Furthermore, TGIF1 and TGIF2 were also present in transcription factors binding to JICD1 associated and activated transcription DNA motifs in the *Sox2* promoter (Supplementary Fig. 9b). TGIFs function as transcriptional repressors by recruiting co-repressors, including HDAC, Sin3a, and CtBP^38, 39, 40^. TGIFs inhibit the expression of TGF-β target genes via interaction with the SMAD complex^41^. Our results revealed that the mRNA level of *Sox2* increased in TGIF2 depleted JICD1-overexpressing cells (Fig. 5c), indicating that TGIF2 acts as a transcriptional repressor under these conditions. To determine whether TGIF2 forms a complex with JICD1 and its role in the JICD1 transcription complex, we first examined the interaction between JICD1 and TGIF2. Under the harsh washing conditions in the co-immunoprecipitation (Co-IP) experiment, the association between DDX17 and JICD1 was previously confirmed and the interaction of JICD1 with TGIF1 or TGIF2 was not detected (Fig. 6a). However, under mild washing conditions, JICD1 interacted with TGIF2, not TGIF1 (Supplementary Fig. 9e), indicating that JICD1 and TGIF2 do not directly interact, but are associated in cooperation with other proteins. Indeed, we observed that TGIF2 was associated with SMAD3 rather than that with DDX17 among JICD1 transcription complex components (Fig. 6b).

**Fig. 6:**
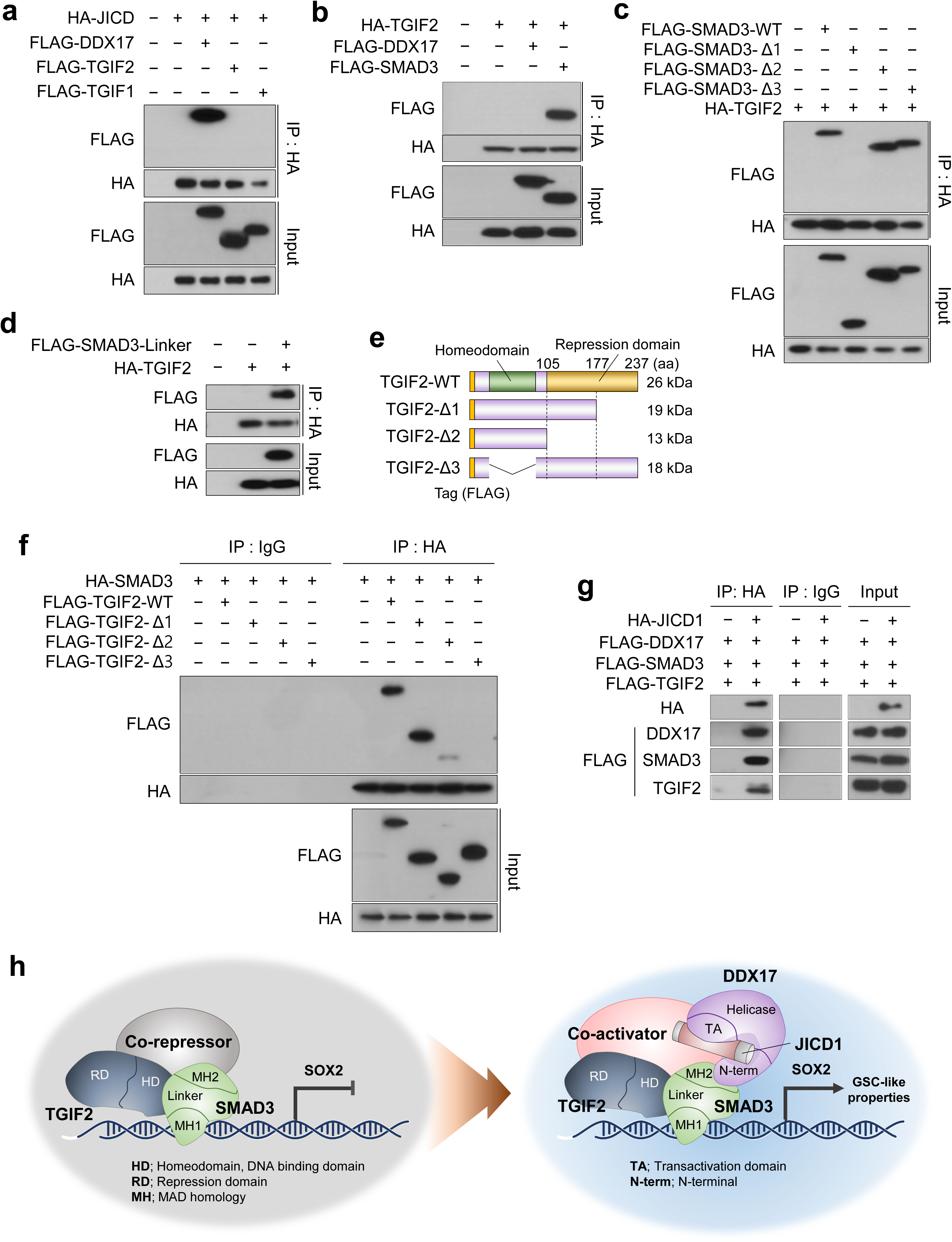
TGIF2 is a binding partner of JICD1/DDX17/SMAD3 complex. **a** Co-IP analysis between FLAG-TGIF1/TGIF2 and HA-JICD1. The association between FLAG-DDX17 and HA-JICD1 was used as a positive control. **b** The association between HA-TGIF2 and FLAG-DDX17/SMAD3 in HEK293T cells. **c** The association between HA-TGIF2 and FLAG-SMAD3 mutants in HEK293T cells. **d** The association between HA-TGIF2 and FLAG-SMAD3 linker domain fragments in HEK293T cells. **e** Schematic diagrams showing TGIF2 mutants. (WT: Wild-type, △: Mutant). **f** The association between HA-SMAD3 and FLAG-TGIF2 mutants in HEK293T cells. **g** The association of JICD1 complex protein, DDX17, SMAD3, and TGIF2 in HEK293T cells. The association between JICD1 and TGIF2 when washed under mild conditions. **h** Schematic diagram that represents the proposed mechanism through which JICD1 forms a transcription complex with DDX17, SMAD3 and TGIF2 and regulates GSC properties via SOX2 expression. See also Supplementary Fig. 9, 10.

Among SMAD3 mutants, TGIF2 interacted with SMAD3 mutants containing the linker domain, and the MH1 domain of SMAD3 further strengthened their interaction (Fig. 6c). Next, we established a SMAD3 mutant consisting of only the linker domain (Fig. 5i) and confirmed its interactions with TGIF2 (Fig. 6d). Therefore, it is plausible that the MH1 domain might stabilize the interaction between SMAD3 and TGIF2.

TGIFs bind to DNA through the homeodomain (HD) and interact with co-repressors through the repression domain (RD)^40^. The interaction between SMAD2 and TGIF1 has been reported^41^, but there is no previous study reporting an interaction between SMAD3 and TGIF2. To gain further insight into their interaction, we constructed various deletion mutations of TGIF2 based on its functional domains (Fig. 6e). RD deleted mutants of TGIF2 did not influence the interaction with SMAD3, however, the HD-deleted mutant failed to interact with SMAD3 (Fig. 6f). However, on partially deleting RD, the interaction with SMAD3 was stronger than that when the entire RD is deleted. Thus, it is likely that a part of the RD domain might stabilize the interaction with SMAD3.

Finally, to confirm the formation of the JICD1 transcription complex, HA-JICD1, FLAG-DDX17, FLAG-SMAD3, and FLAG-TGIF2 were co-transfected into HEK293T cells. As determined by immunoprecipitation with anti-HA, JICD1 was associated with DDX17, SMAD3, and TGIF2 (Fig. 6g). Overall, JICD1 formed a transcription complex with DDX17, SMAD3, and TGIF2 (Fig. 6h and Supplementary Fig. 10). This JICD1 transcriptional complex promotes tumorigenicity with cancer stemness by increasing SOX2 expression, suggesting that JICD1 is an important target in eliminating cancer stem cells.

## Discussion

NICD1 has various functional domains, including the PEST domain, transactivation domain, nuclear localization signal domain, ankyrin repeats (ANK) domain, and RBPJ-associated molecule (RAM), however, JICD1 only has a functional domain called the PDZL domain^42^. NICD1 interacts with CSL through the RAM and ANK domains, as well as Mastermind through the ANK domain^43, 44^. JICD1 is extremely short when compared with NICD1, and hence its interaction with other proteins is limited. However, in the present study, we demonstrated that JICD1 acts as a transcriptional cofactor in the formation of the JICD1/DDX17/SMAD3/TGIF2 complex.

We identified JICD1-binding proteins in the nucleus and cytoplasm using affinity purification and tandem mass spectrometry. Among these proteins, DDX17 functions as a transcriptional activator of SOX2 in cooperation with JICD1. DDX17 reportedly forms a homodimer or heterodimer with DDX5^35^. The core helicase domains of DDX17 and DDX5 are 90% homologous, but the N-terminal and C-terminal display only 60% and 30% homology, respectively. Both proteins bind to the same transcription factors, including β-catenin, nuclear factor of activated T-cells 5, estrogen receptor alpha, and androgen receptor^32, 35^. However, in most cases, the transcriptional regulation of DDX17 and DDX5 is not redundant and depends on binding factors at their N-terminal and C-terminal. Indeed, JICD1 was bound to the N-terminal and C-terminal of DDX17, but not to DDX5 in our study (data not shown).

A recent report has suggested that DDX17 maintains long-term chromatin conformation by regulating DNA looping and converting tumor cells into an invasive phenotype by modulating a chromatin-binding factor^45, 46^. Furthermore, SMADs regulate gene expression by recruiting chromatin modifiers and remodelers, including histone acetyltransferase p300, ATP-dependent helicase SMARCA4, H3K27 demethylase JMJD3, and mediator components^47^. Incidentally, the JICD1-upregulated DEGs included genes associated with chromatin remodeling (Supplementary Fig. 3a), which is consistent with the presence of the chromatin-modulating CCCTC-binding factor within the JICD1-binding DNA motif. Therefore, the JICD1 transcription complex may not only directly regulate transcription but also alter the chromatin structure.

SMAD3 is one of the main signal transducers for TGFβ receptors^48^. SMADs consist of three subtypes: receptor-regulated SMADs (R-SMADs), common partner SMAD (co-SMAD), and inhibitory SMADs (I-SMADs). In the canonical pathway, two R-SMADs and one co-SMAD form a trimer, which is a predominant effector regulating transcription. We confirmed that the JICD1/DDX17 complex binds to DNA via the transcription factor SMAD3. Furthermore, SMAD3- and SMAD4-binding DNA motifs exist in the JICD1-upregulated DEG promoter (Supplementary Fig. 9d). Based on these results, JAG1, independent of Notch receptor signaling, can crosstalk with the TGFβ signal via the JICD1 transcription complex.

Interestingly, TGIF2, known as a co-repressor that regulates the transcription of TGFβ-responsive genes by interacting with SMADs, interacted with the JICD1 transcription complex (Fig. 6). It remains unclear whether TGIF2 acts as a repressor in the JICD1 transcription complex. The linker domain of R-SMADs is phosphorylated by CDK8 and CDK9 in the nucleus, and this phosphorylation regulates interaction with co-activators or co-repressors^49^. In the JICD1 transcription complex, SMAD3 interacts with TGIF2 through the linker domain (Fig. 6c, d). Therefore, additional investigations are needed to determine whether SMAD3 phosphorylation could alter the associated co-activators and co-repressors and assess whether the activity of the cofactor could be altered depending on the phosphorylation status of SMAD3.

The Notch signaling pathway delivers unidirectional signaling from the Notch ligand to the Notch receptor. In contrast, our results revealed that cells expressing the JAG1 ligand generated the JICD1 transcription complex. JICD1 and DDX17 may act as a switch for the transcription complex, using SMAD3 as a molecular hub to recruit chromatin modifiers and remodelers. These complexes regulate the acetylation of the *Sox2* promoter, and increased SOX2 induces oncogenic functions such as stemness, tumorigenesis, invasion, and resistance to apoptosis. Thus, understanding the precise mechanism of JAG1 and JICD1 in oncogenic transformation provides a new strategy to overcome the limitations of current anticancer drugs targeting the Notch receptor signaling pathway.

## Methods

### Cell lines and culture conditions

Ink4a/Arf^-/-^ astrocytes were isolated from murine cerebral cortices of 5-day-old Ink4a/Arf^-/-^ knockout mice^27^. The human glioma cell line LN18 (RRID: CVCL_0392) was purchased from American Type Culture Collection (ATCC, Manassas, VA, USA). Ink4a/Arf^-/-^ astrocytes and LN18 were cultured in Dulbecco’s Modified Eagle Medium (DMEM, Cat. #17-605E; Lonza, Basel, Switzerland) supplemented with 10% fetal bovine serum (FBS, Cat. #7101; Biotechnics Research, Inc., Canton, OH, USA), 1% penicillin and streptomycin (P/S, Cat. #17-602E; Lonza), 1% L-glutamine (Cat. #17-605E; Lonza), and gentamicin sulfate (50 μg/mL, Cat. #MT-61098-RF; Mediatech, Mooresville, NC, USA). GSCs were obtained from Dr. Ichiro Nakano (The Ohio State University, Columbus, OH, USA)^50^. GSCs were cultured in DMEM and Ham’s F-12 (DMEM/F12, Cat. #12-719Q; Lonza) supplemented with 1% P/S (Cat. #17-602E; Lonza), 1% L-glutamine, 0.2% B27 (Cat. #17504-044; Invitrogen, Carlsbad, CA, USA), epidermal growth factor (EGF, 20 ng/mL, Cat. #236-EG; R&D Systems, Minneapolis, MN, USA), basic fibroblast growth factor (bFGF, 20 ng/mL, Cat. #4114-TC; R&D Systems), and gentamicin sulfate (50 μg/mL).

### Plasmids and gene transduction

Human JAG1, NICD1, HA-JICD1, HA-SMAD3, FLAG-SMAD3, HA-DDX17, FLAG-TGIF1, HA-TGIF2, and FLAG-TGIF2 were cloned into the pcDNA3.1-puro vector. The pcDNA3.1-FLAG-DDX17-neo was kindly provided by Dr. Didier Auboeuf (Centre Léon Bérard, Lyon, France)^45^. The JICD1, DDX17, SMAD3, and TGIF2 mutants were amplified by PCR and then cloned into the pcDNA3.1-puro or pcDNA3.1-neo vector. Short hairpin RNAs (shRNAs) targeting mouse Ddx17, Smad3, and Tgif2 were purchased from Sigma-Aldrich (shDdx17-4: TRCN0000071082, shDdx17-5: TRCN0000312878, shSmad3-2: TRCN0000089023, shSmad3-3: TRCN0000089025, shTgif2-1: TRCN0000075523). shTgif1 was kindly provided by Dr. Jonghwan Kim (University of Texas at Austin, Austin, TX, USA).

To construct knockdown cell lines, each shRNA vector was transfected with second-generation lentiviral packaging plasmids △8.9 and pVSV.G using the PolyExpress^TM^ in vitro DNA transfection reagent (Cat. #EG1072; Excellgen, Rockville, MD, USA) into transformed human embryonic kidney 293 cells, HEK293T (Cat. #CRL-3216, ATCC). Twenty-four hours after transfection, the culture medium was harvested. The lentivirus was filtered through a 0.45 μm syringe filter and concentrated with Lenti-X^TM^ Concentrator (Cat. #631231; Clontech, Mountain View, CA, USA). The cells were infected with the viruses in the presence of 6 μg/mL polybrene (Cat. #107689; Sigma-Aldrich, St. Louis, MO, USA). The overexpressing vectors were transfected into Ink4a/Arf^-/-^ astrocytes using the Neon^TM^ Transfection System 10μl Kit (Cat. #MPK1096; Invitrogen) according to the manufacturer’s instructions.

### Quantitative reverse transcription-polymerase chain reaction (qRT-PCR)

Total RNA was isolated from cells using QIAzol lysis reagent (Cat. #79306; Qiagen, Valencia, CA, USA) according to the manufacturer’s instructions. RNase-free, DNase-treated RNA (1 μg) was used as the template to synthesize cDNA using the RevertAid First-Strand cDNA Synthesis Kit (Cat. #K1621; ThermoFisher Scientific, Waltham, MA, USA). qRT-PCR analysis was performed on an iCycler IQ real-time detection system (Bio-Rad, Hercules, California, USA) using the IQ Supermix with SYBR-Green (Cat. #RR420A; TaKaRa Bio, Shiga, Japan). Gene expression was quantified using the standard 2^−ΔΔCt^ method as described previously. The expression levels of target genes were normalized to that of GAPDH.

### Co-IP and western blot analysis

For the Co-IP experiment, HEK293T cells were co-transfected with the previously described plasmids and the proteins were extracted using the Pierce^TM^ IP lysis buffer (Cat. #87787; Invitrogen) supplemented with 1 mM phenylmethylsulfonyl fluoride (PMSF, Cat. #10837091001; Sigma-Aldrich), protease inhibitor (Cat. #11836-153-001; Roche, Basel, Switzerland) and phosphatase inhibitors (1 mM NaF, 1 mM Na3VO4, 1 mM β-glycerophosphate, and 2.5 mM sodium pyrophosphate). The lysates were pre-cleared with Protein A/G agarose (Cat. #20333 and #20399; Thermo Fisher Scientific), and proteins (0.5-1 mg) were precipitated using the anti-FLAG antibody (1:100, Cat. #F7425; Sigma-Aldrich), anti-HA antibody (1:100, Cat. #3724s; Cell Signaling Technology, Beverly, MA, USA) and Protein A/G agarose. The FLAG- or HA-binding proteins were washed with IP buffer and eluted with NuPAGE^TM^ LDS sample buffer (Cat. #NP0007; Thermo Fisher Scientific) for 3-5 min at 100°C. The eluted proteins were separated by SDS-polyacrylamide gel electrophoresis (PAGE) and transferred to polyvinylidene fluoride (PVDF) membranes (Cat. #BSP0161; Pall Corporation, New York, NY, USA). The membranes were blocked with 5% non-fat milk and incubated with the following antibodies: anti-FLAG (1:1000, Cat. #F1804; Sigma-Aldrich), and anti-HA (1:1000, Cat. #H9658; Sigma-Aldrich). Following washing, the membranes were incubated with a horseradish peroxidase-conjugated anti-IgG secondary antibody (Pierce, Rockford, IL, USA) and visualized with SuperSignal West Pico Chemiluminescent Substrate (Cat. #EBP-1073; ELPIS Biotech, Seoul, South Korea).

For western blotting, whole-cell extracts were prepared with radioimmunoprecipitation assay (RIPA) lysis buffer (Cat. #CBR002; LPS solution, Daejeon, South Korea) comprised of 150 mM NaCl, 1% NP-40, 0.1% SDS, and 50 mM Tris (pH 7.4) containing 1 mM NaF, 1 mM Na3VO4, 1 mM β-glycerophosphate, 2.5 mM sodium pyrophosphate, and protease inhibitor (Cat. #11836-153-001; Roche). Nuclear and cytosolic proteins were prepared using a nuclear/cytosol fractionation kit (Cat. #K266100; BioVision, Milpitas, CA, USA), according to the manufacturer’s instructions. The cytosolic and nuclear extractions were used for western blot or tandem affinity purification analysis. The proteins were quantified using the Bradford assay reagent (Cat. #500-0006; Bio-Rad) according to the manufacturer’s instructions. Protein (10-50 μg) was separated by 8% SDS-PAGE and immunoblotting was performed as described above. The primary antibodies were used with the following antibodies: anti-JAG1 (1:500, Cat. #sc-6011; Santa Cruz Biotechnology, Dallas, TX, USA), anti-HA (1:1000, Cat. #3724s; Cell Signaling Technology), anti-LAMIN B2 (1:500, Cat. #sc-6216; Santa Cruz Biotechnology), anti-α-tubulin (1:10000, Cat. #T9026; Sigma-Aldrich), anti-cleaved Notch1 (1:1000, Cat. #2421L; Cell Signaling Technology), anti-SOX2 (1:1000, Cat. #AF2018; R&D Systems), anti-DDX17 (1:1000, Cat. #sc-27112; Santa Cruz Biotechnology), anti-SMAD3 (1:500, Cat. #ab28379; Abcam, Cambridge, UK). α-Tubulin was used as a loading control.

### Cell viability assay

In brief, cells were seeded in 96-well plates (5 × 10^2^ cells/well). After 24 h, cells were treated with BCNU (Cat. #C0400; Sigma-Aldrich) or TMZ (Cat. # T2577; Sigma-Aldrich) for 48 h and then assayed for cell viability. Irradiation was conducted with defined doses of ^137^Cs gamma-ray irradiation using an IBL 473C irradiator (CIS Bio-International, Codolet, France). The irradiated cells were seeded in 96-well plates (5 × 10^2^ cells/well), and cell viability was evaluated after 48 h. Cell viability was determined using the 4-[3-(4-iodophenyl)-2-(4-nitrophenyl)-2H-5tetrazolio]-1,3-benzene disulfonate (WST)-based assay (CytoX, Cat. #CYT3000; LPS solution), according to the manufacturer’s instructions. CytoX reagent was added to each well, and cells were incubated at 37℃ for 2 h. The relative viability of cells was determined by measuring the absorbance at 450 nm using a PowerWaveXS microplate-reader (Bio-Tek Instruments, Inc., VT, USA). Experiments were repeated three times, and data represent the mean of replicate wells ± standard error of the mean (S.E.M.).

### Immunofluorescence

HEK293T cells were transfected with plasmids for 48 h and fixed with 4% paraformaldehyde for 20 min at 25℃, followed by permeabilization with PBS plus 0.3% Triton X100, 10% serum and 0.1% bovine serum albumin (BSA). The cells were stained with JAG1 antibody overnight at 4℃. Then, the cells were washed with ice-cold PBS three times for 5 min each and incubated with Alexa 594 conjugated secondary antibody at 25℃ for 2 h. After staining with DAPI (1:1,000, Cat. #D9542; Sigma-Aldrich) for 5 min, slides were mounted in mounting solution (Cat. #P36930; Invitrogen) and stored at 4°C in the dark. Fluorescence was detected using a confocal laser-scanning microscope (LSM700 and LSM800; Carl Zeiss, Jena, Germany).

### Intracranial xenograft model

All mouse experiments were approved by the Animal Care Committee of Korea University according to the government and institutional guidelines and regulations of the Republic of Korea. Ink4a/Arf^-/-^ astrocytes were harvested by trypsinization and washed with phosphate-buffered saline (PBS). Cell viability was determined by the trypan blue exclusion method. Single-cell suspensions with >90% viability were used for *in vivo* experiments. Cells (1 × 10^5^ cells/3μL PBS) were stereotactically injected into the left striatum of 5-week-old BALB/c nu/nu mice (coordinates relative to the bregma: anterior-posterior +2 mm, medial-lateral +2 mm, and dorsal-ventral -3 mm). To compare the tumor histology, all mice were simultaneously sacrificed when the first mouse showed neurological symptoms.

### Hematoxylin and eosin staining and immunohistochemistry

Tumor-bearing mice were perfused with PBS and 4% paraformaldehyde. Intracranial tumor tissues were embedded in paraffin, sectioned (4 µ m in thickness), and placed on glass slides. After deparaffinization and hydration, the tissue slides were treated with hematoxylin (Cat. #105174; Merck, Damstadt, Germany) for 5 min and rinsed with tap water. All slides were incubated in eosin solution (Cat. #109844; Merck) for 30 s and then washed with distilled water.

For immunohistochemistry, the patient tumor tissue slides were stained with anti-JAG1 (Cat. #sc-6011; Santa Cruz Biotechnology) for 12 h at 4°C. After washing, the slides were incubated with a biotin-conjugated secondary antibody for 1 h and with VECTASTAIN ABC Reagent (Cat. #PK- 4000; Vector Laboratories, Burlingame, CA, USA) for 30 min at 20°C. The slides were then incubated in a peroxidase substrate solution (Cat. #PK-4100; Vector Laboratories) and counterstained with hematoxylin. Cell imaging was performed using an Epifluorescence and Brightfield microscope (Axioimager M1; Carl Zeiss).

### *In vitro* invasion assay

An invasion assay was performed using 24-well Transwell units (Cat. #3422; Corning Coster, New York, NY, USA) coated with 1mg/mL Matrigel (Cat. #BD354234; BD Bioscience, San Jose, CA, USA), which was allowed to settle for 4 h at 37℃. Next, 1.0×10^5^ cells suspended in 100 μL serum-free DMEM were seeded into the upper chamber, and 500 μL serum-containing DMEM was added to the lower chamber. The cells invaded for 48 h at 37℃. Next, non-migrated cells from the upper chamber were removed, and migrated cells on the lower side of the membrane were fixed, stained with crystal violet, and dried. Images of the entire upper chamber were obtained, and the invasion area was quantified using ImageJ (https://imagej.nih.gov).

### *In vitro* limiting dilution sphere formation assay

For an *in vitro* limiting dilution assay, decreasing numbers of cells (20, 10, 5, 2, and 1) were manually seeded in individual wells in 96-well plates containing DMEM/F12 with B27, EGF, and bFGF (n = 18). Neurospheres larger than 10 µ m in diameter were counted by light microscopy after 14 days. Stem cell frequency was calculated using the ELDA software, available at http://bioinf.wehi.edu.au/software/elda.

### Tandem affinity purification

HA-beads (Cat. #E6779; Sigma-Aldrich) were washed twice with NETN buffer (0.5% NP-40, 20mM Tris-Cl pH8.0, 100 mM NaCl, 1.5 mM MgCl2, and 1mM EDTA). Them, the washed HA-beads were added to the cytosolic and nuclear fractions, and mixed by rotation for 3 h at 4°C. The supernatant was removed and washed twice with NETN buffer and PBS. The HA-binding proteins were eluted with NuPAGE^TM^ LDS sample buffer and resolved by SDS-PAGE. Coomassie-stained bands of SDS-PAGE were analyzed by mass spectrometry.

### Chromatin Immunoprecipitation (ChIP) assay

For crosslinking, the cells were treated with formaldehyde (final concentration 1%) in the culture medium for 10 min at 20°C. The reaction was quenched with glycine (final concentration 0.125M) for 5 min at 20°C. Then, the cells were washed twice with cold PBS and harvested using a cell scraper. The cell pellet was resuspended in lysis buffer (0.5% NP40, 85 mM KCL, 5 mM Pipes, 1 mM PMSF, 0.01 mg/mL aprotinin, and 0.01 mg/mL leupeptin). After incubation for 10 min on ice, the supernatant was removed by centrifugation (5,000 rpm, 5 min), and the pellet was lysed with nuclear lysis buffer (1% SDS, 10mM EDTA, 50 mM Tris pH 8.1, 1 mM PMSF, 0.01 mg/mL aprotinin, and 0.01 mg/mL leupeptin) for 10 min on ice. Cross-linked chromatin was fragmented to the 0.3–1.5 kb range by focused-ultrasonication. The insoluble fraction was removed by centrifugation and the supernatant was pre-cleared with Protein A/G/sperm DNA beads for 2 h at 4°C. For ChIP, 300 µ g of chromatin DNA was incubated with anti-HA (Cat. #3724s; Cell Signaling Technology), anti-H3K9Ac (Cat. #07-352; EMD Millipore, Billerica, MA, USA), at 4°C overnight, followed by 60 µ L of Protein A/G/sperm DNA beads for 2 h at 4°C. The beads were then washed and eluted. Chromatin crosslinking was reversed by adding 5 M NaCl at 65°C overnight. Residual RNA was digested with RNase A for 30 min at 37°C, and the DNA was purified using the DNA purification kit (Cat. #EBP-1004; ELPIS Biotech).

### ChIP-seq analysis

Isolated cross-linked chromatin samples were pre-cleared with Protein A/G without sperm DNA. Libraries for ChIP-seq were constructed using NEBNext® Ultra^TM^ DNA Library Prep Kit for Illumina (Cat. #E7370; New England Biolabs, Ipswich, MA, USA), according to the manufacturer’s instructions. Pre-cleared ChIP DNA was ligated with adaptors. After purification, PCR was performed with adaptor-ligated DNA and index primers for multiplex sequencing. The library was purified using magnetic beads to remove reaction components, and the size of library fragments was assessed by Agilent 2100 bioanalyzer (Agilent Technologies, Santa Clara, CA, USA). High-throughput sequencing was performed as single-end 75 sequencing using NextSeq 500 (Illumina, San Diego, California, USA). The reads were trimmed and aligned using Bowtie2. Bowtie2 indices were either generated from the genome assembly sequence or the representative transcript sequences for aligning to the genome and transcriptome. We used Model-based Analysis of ChIP-seq (MACS) to identify peaks from the alignment file. Gene classification of the hits was performed using DAVID (http://david.abcc.ncifcrf.gov/) and Medline databases (http://www.ncbi.nlm.nih.gov/). We performed motif analysis on the promoter region (-10 kb) of the JICD1-upregulated DEGs (n = 536) within the ChIP-sequencing results. FindMotifBySymbols in R were used to search for motifs present in the peaks from which we obtained the results of motifs that match the consensus sequence in the promoter region.

### Luciferase reporter assay

To compare the transcriptional activity to CSL sequences, the pGL3-CSL WT-control vector and pRL-TK were transiently co-transfected into control, HA-JICD1, and NICD1 overexpressing Ink4a/Arf^-/-^ astrocytes using the PolyExpress^TM^ in vitro DNA transfection reagent. Promoter activity was analyzed using Dual-Glo Luciferase Assay System (Cat. #E2920; Promega, Madison, WI, USA). Transfection efficiency was normalized to the activity of Renilla luciferase according to the manufacturer’s instructions.

### RNA-sequencing and *in silico* analysis

RNA-Seq was performed at the Beijing Genomics Institute (BGI). DEGs were calculated with reads per kilobase per million (RPKM) values of the two biological replicates and grouped based on >2-fold change in expression. GO analysis was performed using DAVID Bioinformatics Resources 6.8 (http://david.ncifcrf.gov/home.jsp)^28^. The RNA-seq data are available from GEO accession GSE126855.

Gene expression and patient prognosis were analyzed in microarray datasets from the REpository for Molecular BRAin Neoplasia DaTa (REMBRANDT) database of the National Cancer Institute (https://caintergator.nci.nih.gov/rembrandt/)^20, 51^. The REMBRANDT database was classified based on the expression of JAG1 and NOTCH1 and further based on the JAG1-upregulated DEGs according to the mean ± S.E.M. value of the enrichment score developed using single sample Gene Set Enrichment Analysis (ssGSEA; http://software.broadinstitute.org/cancer/software/genepattern)^52^.

### Quantification and Statistical analysis

All experiments were performed more than three times. Data were analyzed using the two-tailed Student’s *t*-test and are reported as the mean ± S.E.M. For statistical analyses of the patient dataset, a log-rank (Mantel-Cox) test and a two-tailed unpaired *t*-test were used and analyzed using GraphPad Prism 6.0 (GraphPad Software, Inc., La Jolla, CA, USA). The level of statistical significance stated in the text was based on *p* values. \**p*<0.05; \*\**p*<0.01; and \*\*\**p*<0.001 were considered statistically significant.

## Supporting information

extension file

## Acknowledgement

We thank all members of the Cancer Growth Regulation Lab for supportive discussions and technical assistance. We thank Dr. Didier Auboeuf (Centre Léon Bérard, Lyon, France) for providing the DDX17 vector, Dr. Jonghwan Kim (University of Texas at Austin, Austin, TX, USA) for supplying the shTgif1 vector, and Dr. Ichiro Nakano (The Ohio State University, Columbus, OH, USA) for providing GSCs. This study was supported by grants from the National Research Foundation of Korea (NRF) funded by the Ministry of Education (2017R1E1A1A01074205 to H. Kim; 2016R1D1A1B03934783 to S.-C. Kim; 2016R1D1A1B03931941 to S.-H Kim; and 2016M3C7A1913844 to S.H. Kim).

## Author Contributions

H.K., E.-J.K., and J.Y.K. designed the experiments and wrote the manuscript. E.J.K. and J.Y.K. performed most of the experiments. S.W.H. and S.S. conducted the *in vitro* experiments. S.-H.C., M.G.P., N.H., and J.J. generated plasmid construction and engineered cell lines. J.Y.K. and X.J. performed the bioinformatics analysis. K.L., H.J.J, S.J.K., S.J., and K.M. assisted the plasmid preparation. S.-O.K. performed tandem affinity purification under the supervision of S.-C.K. S.-H.K., and S.H.K. provided technical support. All authors discussed the study and commented on the manuscript.

## Reference

1. Wen, P.Y. & Kesari, S. Malignant gliomas in adults. N. Engl. J. Med. 359, 492–507 (2008).

2. Stopschinski, B.E., Beier, C.P., Beier, D. Glioblastoma cancer stem cells--from concept to clinical application. Cancer Lett. 338, 32–40 (2013).

3. Tarasov, V.V. et al. Feasibility of Targeting Glioblastoma Stem Cells: From Concept to Clinical Trials. Curr.Top. Med. Chem. 19, 2974–2984 (2019).

4. Vescovi, A.L., Galli, R., Reynolds, B.A. Brain tumour stem cells. Nat. Rev. Cancer 6, 425–436 (2006).

5. Lobry, C., Oh, P., Mansour, M.R., Look, A.T., Aifantis I. Notch signaling: switching an oncogene to a tumor suppressor. Blood 123, 2451–2459 (2014).

6. Louvi, A., Artavanis-Tsakonas S. Notch signalling in vertebrate neural development. Nat. Rev. Neurosci. 7, 93–102 (2006).

7. Wang, R. et al. iNOS promotes CD24(+)CD133(+) liver cancer stem cell phenotype through a TACE/ADAM17-dependent Notch signaling pathway. Proc. Natl. Acad. Sci. U. S. A. 115, E10127–e10136 (2018).

8. Rajakulendran, N. et al. Wnt and Notch signaling govern self-renewal and differentiation in a subset of human glioblastoma stem cells. Genes Dev. 33, 498–510 (2019).

9. Bray, S.J. Notch signalling in context. Nat. Rev. Mol. Cell Biol. 17, 722–735 (2016).

10. Schroeter, E.H., Kisslinger, J.A., Kopan, R. Notch-1 signalling requires ligand-induced proteolytic release of intracellular domain. Nature 393, 382–386 (1998).

11. Lubman, O.Y., Ilagan, M.X., Kopan, R., Barrick, D. Quantitative dissection of the Notch:CSL interaction: insights into the Notch-mediated transcriptional switch. J. Mol. Biol. 365, 577–589 (2007).

12. Ross, D.A. & Kadesch, T. The notch intracellular domain can function as a coactivator for LEF-1. Mol. Cell. Biol. 21, 7537–7544 (2001).

13. Borggrefe, T. & Oswald, F. The Notch signaling pathway: transcriptional regulation at Notch target genes. Cell. Mol. Life Sci. 66, 1631–1646 (2009).

14. Ntziachristos, P, Lim, J.S., Sage, J., Aifantis, I. From fly wings to targeted cancer therapies: a centennial for notch signaling. Cancer Cell 25, 318–334 (2014).

15. Li, D., Masiero, M., Banham, A.H., Harris, A.L. The notch ligand JAGGED1 as a target for anti-tumor therapy. Front. Oncol. 4, 254 (2014).

16. Qiu, X.X. et al. High Jagged1 expression is associated with poor outcome in primary glioblastoma. Med. Oncol. 32, 341 (2015).

17. Purow, B.W. et al. Expression of Notch-1 and its ligands, Delta-like-1 and Jagged-1, is critical for glioma cell survival and proliferation. Cancer Res. 65, 2353–2363 (2005).

18. LaVoie, M.J. & Selkoe, D.J. The Notch ligands, Jagged and Delta, are sequentially processed by alpha-secretase and presenilin/gamma-secretase and release signaling fragments. J. Biol. Chem. 278, 34427–34437 (2003).

19. He, W., Hu, J., Xia, Y., Yan, R. β-site amyloid precursor protein cleaving enzyme 1(BACE1) regulates Notch signaling by controlling the cleavage of Jagged 1 (Jag1) and Jagged 2 (Jag2) proteins. J. Biol. Chem. 289, 20630–20637 (2014).

20. Madhavan, S., Zenklusen, J.C., Kotliarov, Y., Sahni, H., Fine, H.A., Buetow, K. Rembrandt: helping personalized medicine become a reality through integrative translational research. Mol. Cancer Res. 7, 157–167 (2009).

21. Verhaak, R.G. et al. Integrated genomic analysis identifies clinically relevant subtypes of glioblastoma characterized by abnormalities in PDGFRA, IDH1, EGFR, and NF1. Cancer Cell 17, 98–110 (2010).

22. Guan, X., et al. Molecular subtypes of glioblastoma are relevant to lower grade glioma. PLoS One 9, e91216 (2014).

23. Saito, N. et al. A high Notch pathway activation predicts response to γ secretase inhibitors in proneural subtype of glioma tumor-initiating cells. Stem Cells 32, 301–312 (2014).

24. Benedito, R. et al. The notch ligands Dll4 and Jagged1 have opposing effects on angiogenesis. Cell 137, 1124–1135 (2009).

25. Lu, J. et al. Endothelial cells promote the colorectal cancer stem cell phenotype through a soluble form of Jagged-1. Cancer Cell 23, 171–185 (2013).

26. Pelullo, M. et al. Kras/ADAM17-Dependent Jag1-ICD Reverse Signaling Sustains Colorectal Cancer Progression and Chemoresistance. Cancer Res 79, 5575–5586 (2019).

27. Jeon, H.M. et al. Inhibitor of differentiation 4 drives brain tumor-initiating cell genesis through cyclin E and notch signaling. Genes Dev. 22, 2028–2033 (2008).

28. Huang, D.W., Sherman, B.T., Lempicki, R.A. Systematic and integrative analysis of large gene lists using DAVID bioinformatics resources. Nat. Protoc. 4, 44–57 (2009).

29. Chen, J., Chen, S., Zhuo, L., Zhu, Y., Zheng, H. Regulation of cancer stem cell properties, angiogenesis, and vasculogenic mimicry by miR-450a-5p/SOX2 axis in colorectal cancer. Cell Death Dis. 11, 173 (2020).

30. Oliphant, M.U.J. et al. SIX2 Mediates Late-Stage Metastasis via Direct Regulation of SOX2 and Induction of a Cancer Stem Cell Program. Cancer Res. 79, 720–734 (2019).

31. Jeon, H.M. et al. ID4 imparts chemoresistance and cancer stemness to glioma cells by derepressing miR-9*-mediated suppression of SOX2. Cancer Res. 71, 3410–3421 (2011).

32. Linder, P. & Jankowsky, E. From unwinding to clamping - the DEAD box RNA helicase family. Nat. Rev. Mol. Cell Biol. 12, 505–516 (2011).

33. Fuller-Pace, F.V. DExD/H box RNA helicases: multifunctional proteins with important roles in transcriptional regulation. Nucleic Acids Res. 34, 4206–4215 (2006).

34. Gates, L.A. et al. Acetylation on histone H3 lysine 9 mediates a switch from transcription initiation to elongation. J. Biol. Chem. 292, 14456–14472 (2017).

35. Fuller-Pace, F.V. The DEAD box proteins DDX5 (p68) and DDX17 (p72): multi-tasking transcriptional regulators. Biochim. Biophys. Acta. 1829, 756–763 (2013).

36. Roberts, A.B., Russo, A., Felici, A., Flanders, K.C. Smad3: a key player in pathogenetic mechanisms dependent on TGF-beta. Ann. N. Y. Acad. Sci. 995, 1–10 (2003).

37. Wotton, D., Lo, R.S., Lee, S., Massagué, J. A Smad transcriptional corepressor. Cell 97, 29–39 (1999).

38. Melhuish, T.A., Gallo, C.M., Wotton, D. TGIF2 interacts with histone deacetylase 1 and represses transcription. J. Biol. Chem. 276, 32109–32114 (2001).

39. Wotton, D., Knoepfler, P.S., Laherty, C.D., Eisenman, R.N., Massagué, J. The Smad transcriptional corepressor TGIF recruits mSin3. Cell Growth Differ. 12, 457–463 (2001).

40. Wotton, D., Lo, R.S., Swaby, L.A., Massagué, J. Multiple modes of repression by the Smad transcriptional corepressor TGIF. J. Biol. Chem. 274, 37105–37110 (1999).

41. Guca, E. et al. TGIF1 homeodomain interacts with Smad MH1 domain and represses TGF-β signaling. Nucleic Acids Res. 46, 9220–9235 (2018).

42. Mašek, J. & Andersson, E.R. The developmental biology of genetic Notch disorders. Development 144, 1743–1763 (2017).

43. Contreras-Cornejo, H. et al. The CSL proteins, versatile transcription factors and context dependent corepressors of the notch signaling pathway. Cell Div. 11, 12 (2016).

44. Wilson, J.J. & Kovall, R.A. Crystal structure of the CSL-Notch-Mastermind ternary complex bound to DNA. Cell 124, 985–996 (2006).

45. Dardenne, E. et al. Splicing switch of an epigenetic regulator by RNA helicases promotes tumor-cell invasiveness. Nat. Struct. Mol. Biol. 19, 1139–1146 (2012).

46. Morgan, S.L. et al. Manipulation of nuclear architecture through CRISPR-mediated chromosomal looping. Nat. Commun. 8, 15993 (2017).

47. Coda, D.M. et al. Distinct modes of SMAD2 chromatin binding and remodeling shape the transcriptional response to NODAL/Activin signaling. Elife 6, (2017).

48. Derynck, R., Zhang, Y., Feng, X.H. Smads: transcriptional activators of TGF-beta responses. Cell 95, 737–740 (1998).

49. Shi, Y. & Massagué, J. Mechanisms of TGF-beta signaling from cell membrane to the nucleus. Cell 113, 685–700 (2003).

50. Mao, P. et al. Mesenchymal glioma stem cells are maintained by activated glycolytic metabolism involving aldehyde dehydrogenase 1A3. Proc. Natl. Acad. Sci. U. S. A. 110, 8644–8649 (2013).

51. Bowman, R.L., Wang, Q., Carro, A., Verhaak, R.G., Squatrito, M. GlioVis data portal for visualization and analysis of brain tumor expression datasets. Neuro Oncol. 19, 139–141 (2017).

52. Subramanian, A. et al. Gene set enrichment analysis: a knowledge-based approach for interpreting genome-wide expression profiles. Proc. Natl. Acad. Sci. U. S. A. 102, 15545–15550 (2005).

