## Supplementary material for "JAG1 intracellular domain acts as a transcriptional cofactor that forms an oncogenic transcriptional complex with DDX17/SMAD3/TGIF2": extension file: Supplemental information_Kim et al.pdf

**Contents**

Supplementary Figures 1 to S9

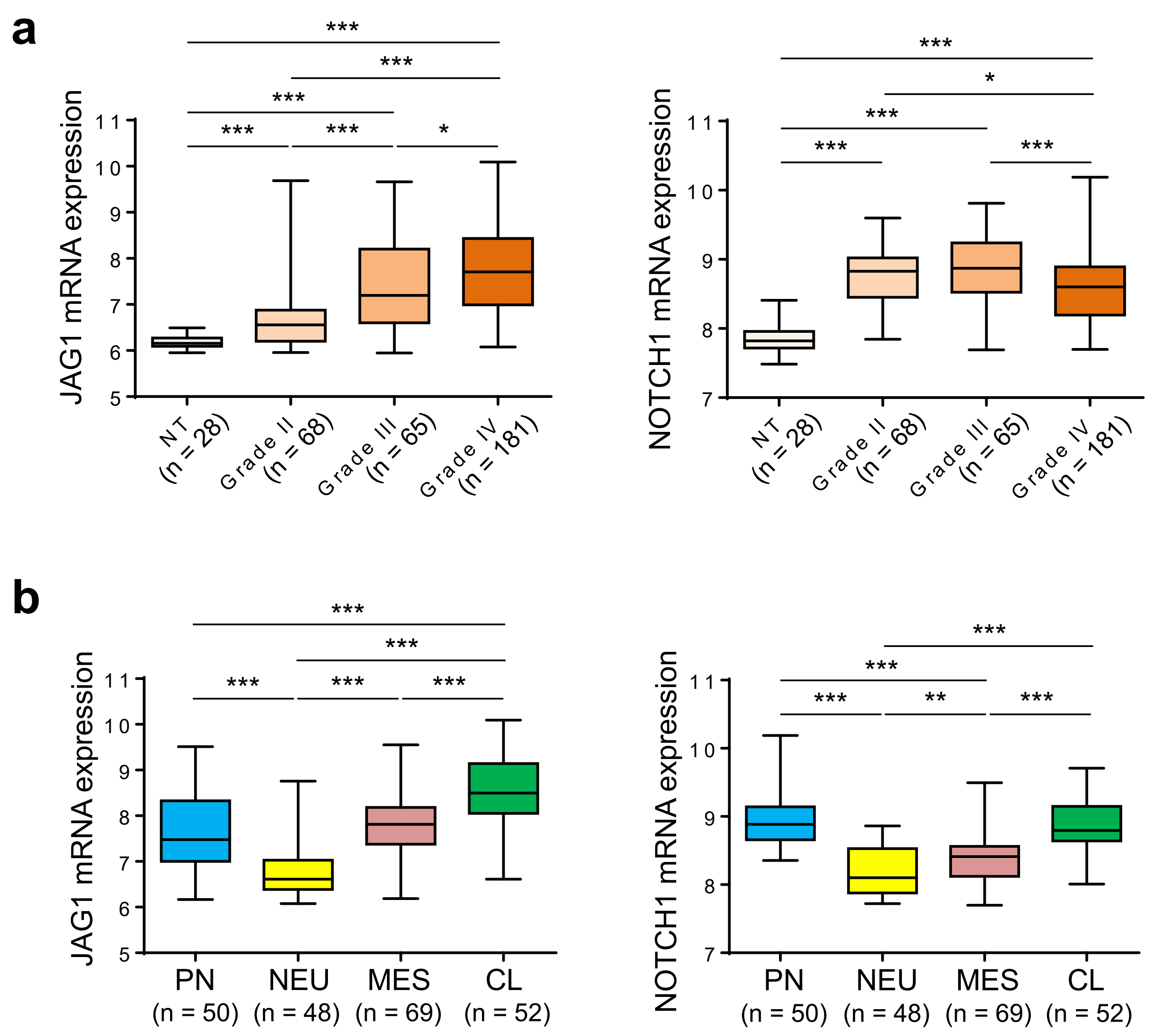


**Supplementary Fig. 1 | The expression patterns of JAG1 and Notch1 according to tumor grades and GBM subtype. Related to Figure 1. a**, Expression levels of JAG1 (left) and NOTCH1 (right) mRNA in non-tumor (NT) WHO grade II, III, and IV from the REMBRANDT database. Unpaired two-tailed t tests: **$p < 0.01$, ***$p < 0.001$. **b**, Expression levels of JAG1 (left) and NOTCH1 (right) mRNA in GBM subtype analyzed using the REMBRANDT database. Proneural (PN), neural (NEU), mesenchymal (MES), classical (CL) subtypes. **$p < 0.01$, ***$p < 0.001$.

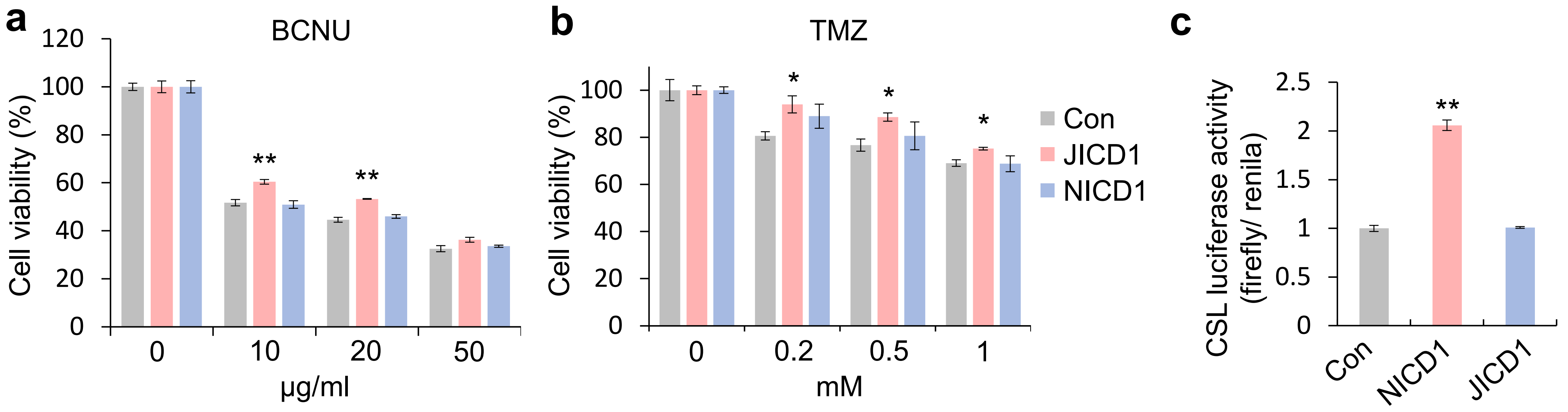


**Supplementary Fig. 2 | JICD1 induces drug resistance and did not demonstrate the transcription activity of the CSL promoter. Related to Figure 2. a**, Drug resistance demonstrated by control, HA-JICD1, and NICD1 overexpressing Ink4a/Arf$^{-/-}$ astrocytes treated with BCNU (0, 10, 20, 50 μM) over 48 h. $^{**}p < 0.01$. **b**, Drug resistance demonstrated by control, HA-JICD1, and NICD1 overexpressing Ink4a/Arf$^{-/-}$ astrocytes treated with TMZ (0, 0.2, 0.5, 1 mM) over 48 h. $^{*}p < 0.05$. **c**, Luciferase reporter analysis to compare transcriptional activity using CSL reporter construct in control, HA-JICD1, and NICD1 overexpressing Ink4a/Arf$^{-/-}$ astrocytes. Relative luciferase activity was normalized to the Firefly/Renilla ratio. $^{**}p < 0.01$.

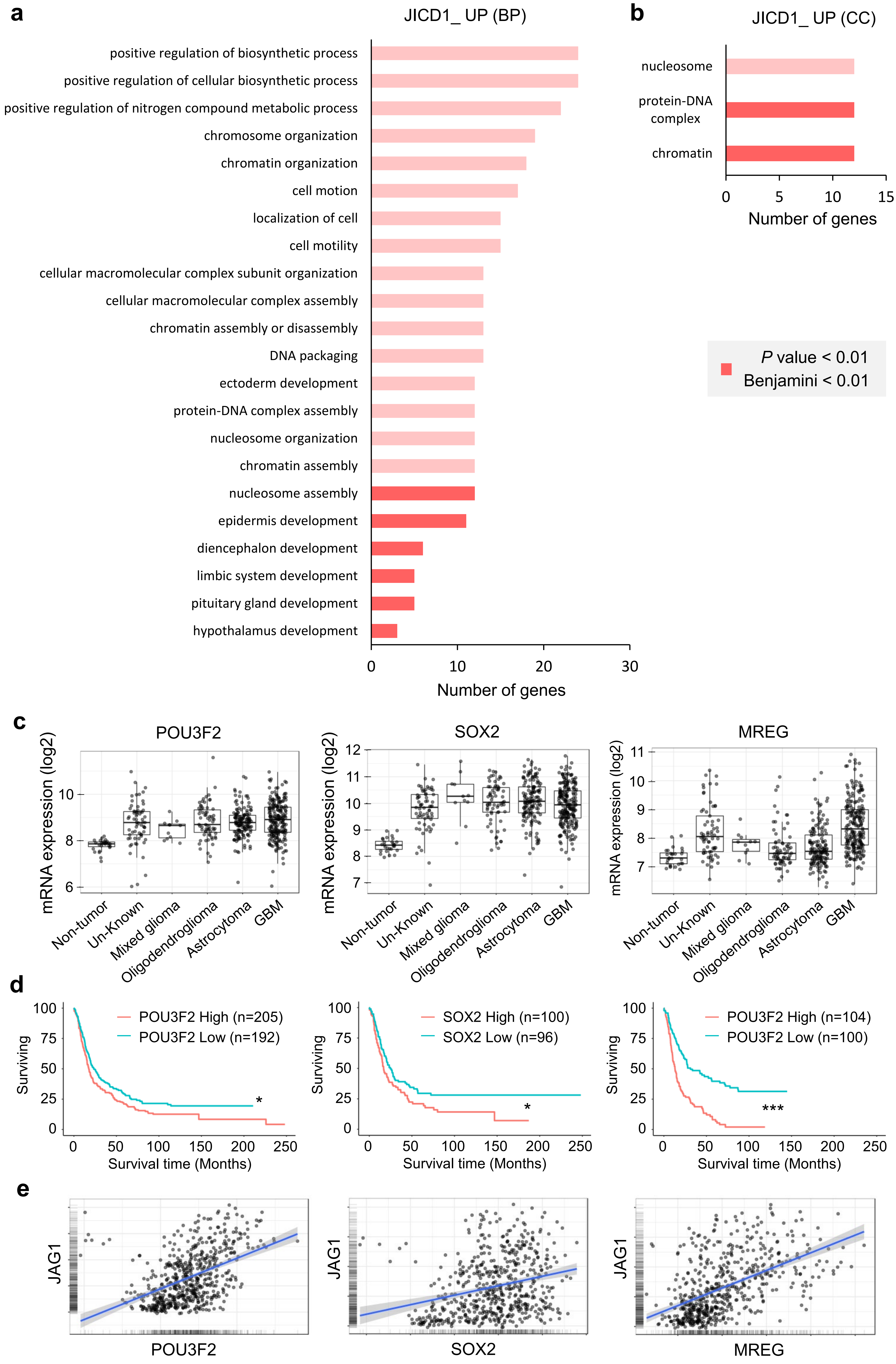


**Supplementary Fig. 3 |** See next page for caption

**Supplementary Fig. 3 | Gene ontology (GO) of JICD1-upregulated DEGs shows chromatin remodeling and development. Related to Figure 3. a**, GO analysis for the biological process (BP) category of JICD1-upregulated (JICD1-UP) DEGs (n=536). **b**, GO analysis for cellular component (CC) category of JICD1-UP DEGs. **c**, The mRNA expression of *Pou3f2, Sox2, and Mreg* was compared according to histology in the REMBRANDT database. **d**, Survival time of patients with glioma according to the expression of *Pou3f2, Sox2, and Mreg*. The patients were divided into two groups (high and low) based on their expression (mean ± S.E.M). *p*-value with a log-rank (Mantel-Cox) test. *$p < 0.05$, ***$p < 0.001$. **e**, The correlation of *JAG1* mRNA with *Pou3f2, Sox2, and Mreg* mRNA in patients with gliomas from the REMBRANDT database.

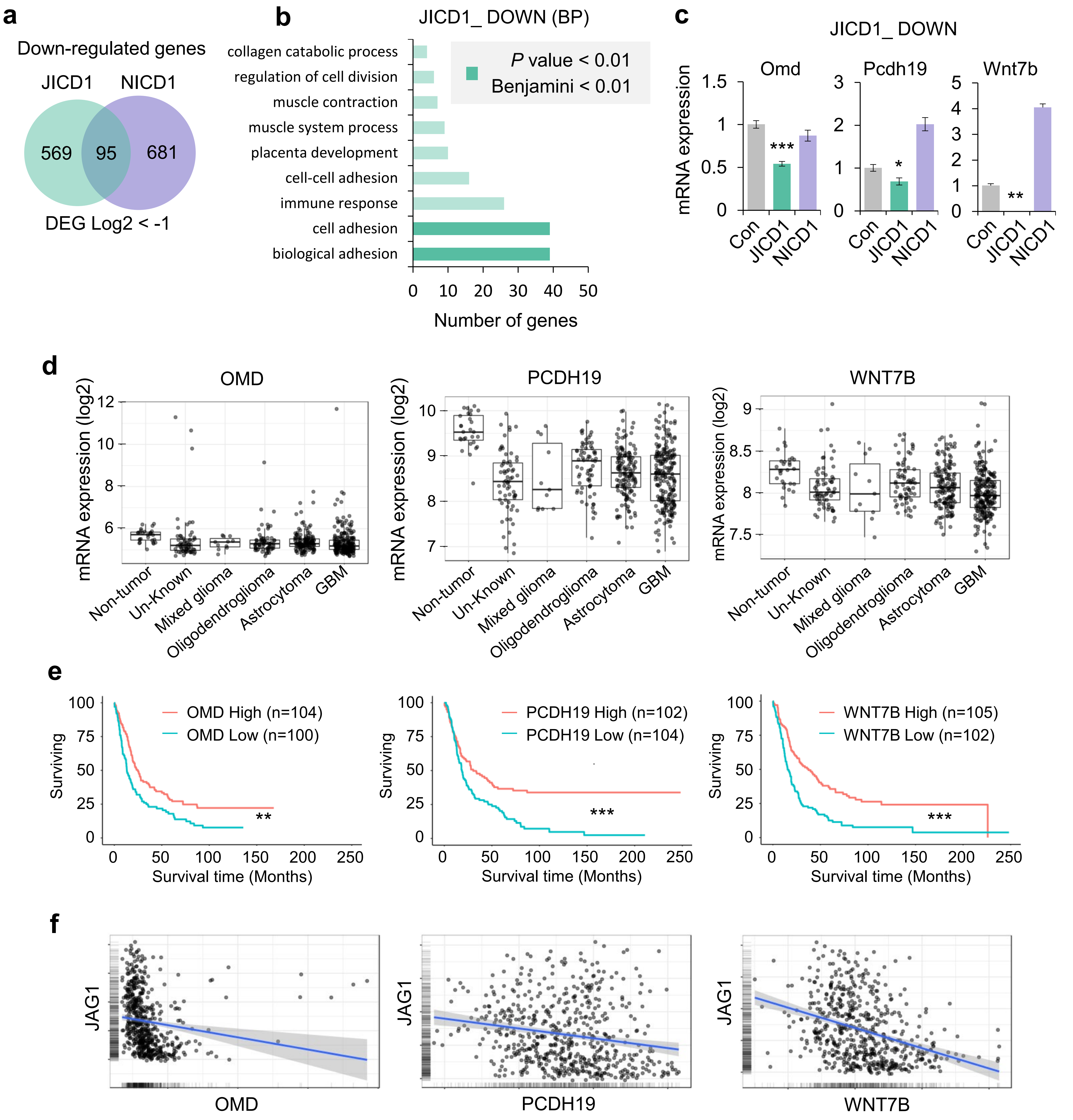


**Supplementary Fig. 4 |** See next page for caption

**Supplementary Fig. 4 | Gene ontology (GO) of JICD1-downregulated DEGs reveals cell adhesion. Related to Figure 3. a**, Unique and overlapping downregulated DEGs in HA-JICD1- and NICD1-overexpressing Ink4a/Arf$^{-/-}$ astrocytes, when compared with control Ink4a/Arf$^{-/-}$ astrocytes. **b**, GO analysis for the BP category of JICD1-downregulated (JICD1-DOWN) DEGs (n=569). **c**, The relative mRNA expression of *Omd*, *Pcdh19*, and *Wnt7b* in control, HA-JICD1- and NICD1-overexpressing Ink4a/Arf$^{-/-}$ astrocytes. *$p < 0.05$, **$p < 0.01$, ***$p < 0.001$. **d**, The mRNA expression of *Omd*, *Pcdh19*, and *Wnt7b* was compared according to histology in the REMBRANDT database. **e**, Survival time of patients with glioma according to the expression of *Omd*, *Pcdh19*, and *Wnt7b*. The patients were divided into two groups (high and low) based on their expression (mean ± S.E.M). *p*-value with a log-rank (Mantel-Cox) test. **$p < 0.01$, ***$p < 0.001$. **f**, The correlation of JAG1 mRNA with *Omd*, *Pcdh19*, and *Wnt7b* mRNA in patients with gliomas from the REMBRANDT database.

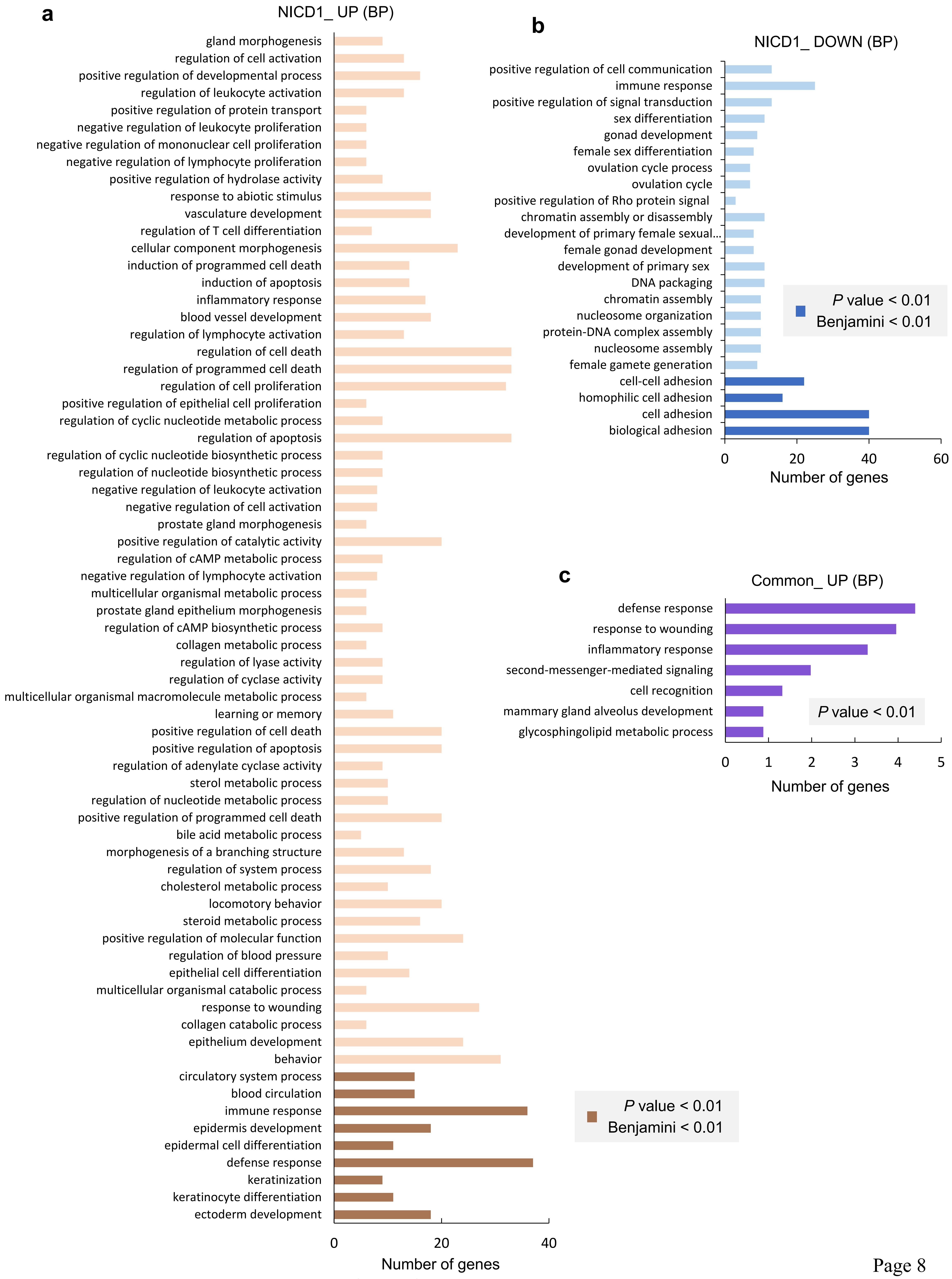


**Supplementary Fig. 5 |** See next page for caption

**Supplementary Fig. 5 | Gene ontology (GO) of NICD1-modulated DEGs showed biosynthetic process, development, immune response, and cell adhesion. Related to Figure 3. a**, GO analysis for the BP category of NICD1-upregulated (NICD1-UP) DEGs (n=803). **b**, GO analysis for the BP category of NICD1-downregulated (NICD1-DOWN) DEGs (n=681). **c**, GO analysis for the BP category of genes upregulated by both JICD1 and NICD1 (Common-UP) DEGs (n=488).

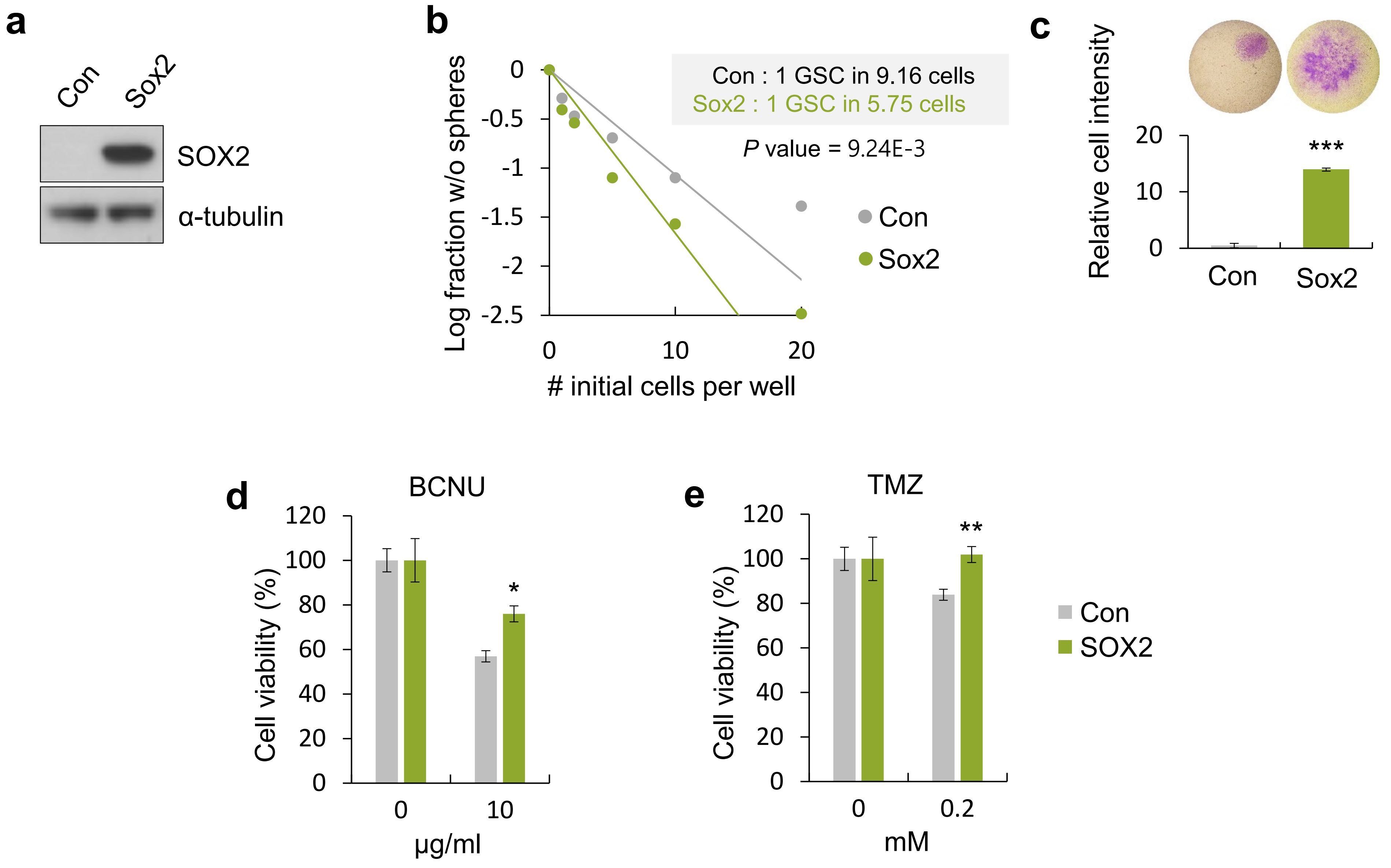


**Supplementary Fig. 6 | SOX2 induces GSC properties. Related to Figure 3. a**, Expression of SOX2 protein in control and SOX2-overexpressing Ink4a/Arf$^{-/-}$ astrocytes. **b**, Sphere forming ability of control, SOX2-overexpressing Ink4a/Arf$^{-/-}$ astrocytes determined performing by an *in vitro* limiting dilution assay. **c**, *In vitro* invasion ability of control, SOX2-overexpressing Ink4a/Arf$^{-/-}$ astrocytes for 48 h. ***$p < 0.001$. **d**, Drug resistance demonstrated by control, SOX2-overexpressing Ink4a/Arf$^{-/-}$ astrocytes treated with BCNU (0, 10 μM) over 48 h. *$p < 0.05$. **e**, Drug resistance demonstrated by control, SOX2-overexpressing Ink4a/Arf$^{-/-}$ astrocytes treated with TMZ (0, 0.2 mM) over 48 h. **$p < 0.01$.

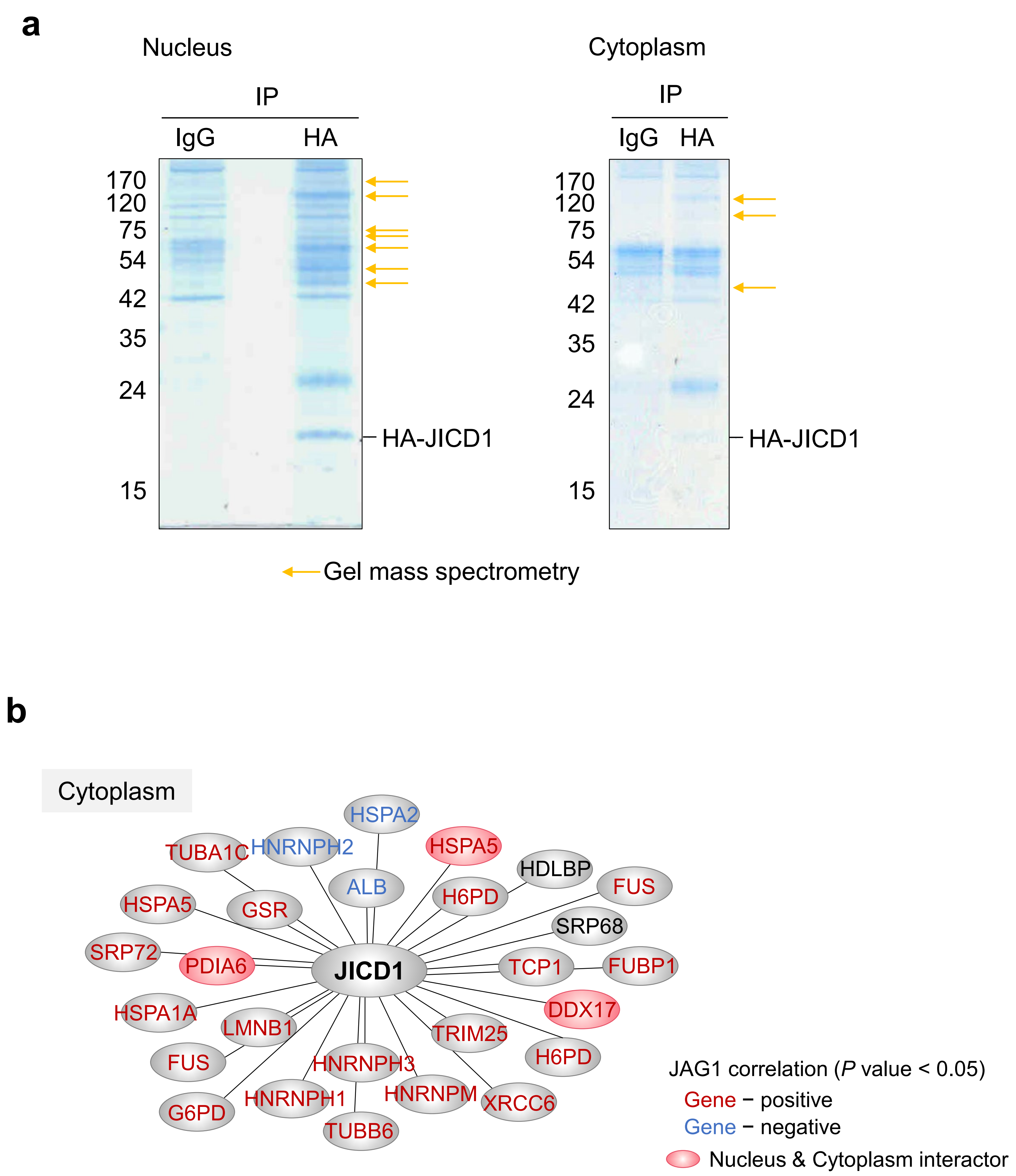


**Supplementary Fig. 7 | JICD1-binding proteins were identified by affinity purification and tandem mass spectrometry. Related to Figure 4. a**, Affinity purified JICD1 binding proteins separated by SDS-PAGE and detected by Coomassie blue staining. Arrows indicate the HA-JICD binding proteins analyzed by gel mass spectrometry. **b**, A schematic diagram that shows JICD1 binding proteins in the cytoplasm. Font color indicates proteins that are correlated with JAG1 in the REMBRANDT database (red: positive, blue: negative, black: no correlation). The red circle indicates proteins that bind to JICD1 in both the nucleus and cytoplasm.

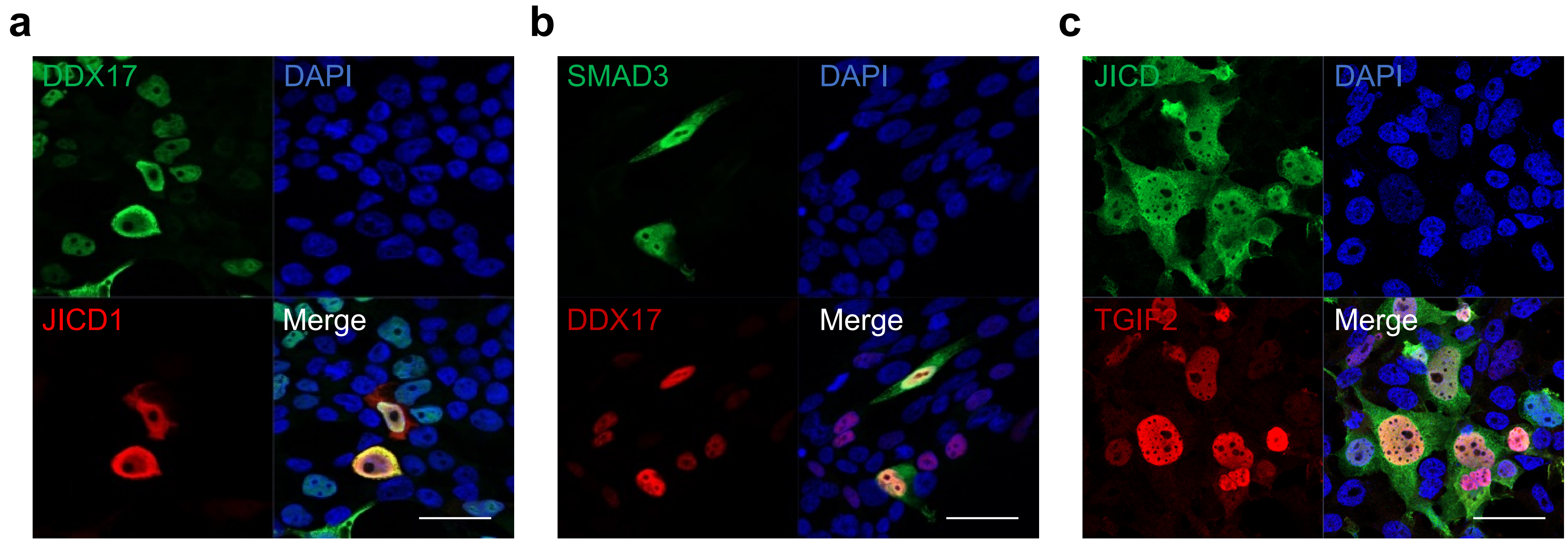


**Supplementary Fig. 8 | JICD1 complex proteins are located at the same location in the cell. Related to Figures 4, 5, and 6. a**, Immunofluorescence showing cellular localization of FLAG-DDX17 (green) and HA-JICD1 (red). DAPI was used to stain the nucleus (blue). Scale bar = 20 μm. **b**, Immunofluorescence showing cellular localization of FLAG-SMAD3 (green) and HA-DDX17 (red). DAPI was used to stain the nucleus (blue). Scale bar = 20 μm. **c**, Immunofluorescence showing cellular localization of HA-JICD1 (green) and FLAG-TGIF2 (red). DAPI was used to stain the nucleus (blue). Scale bar = 20 μm.

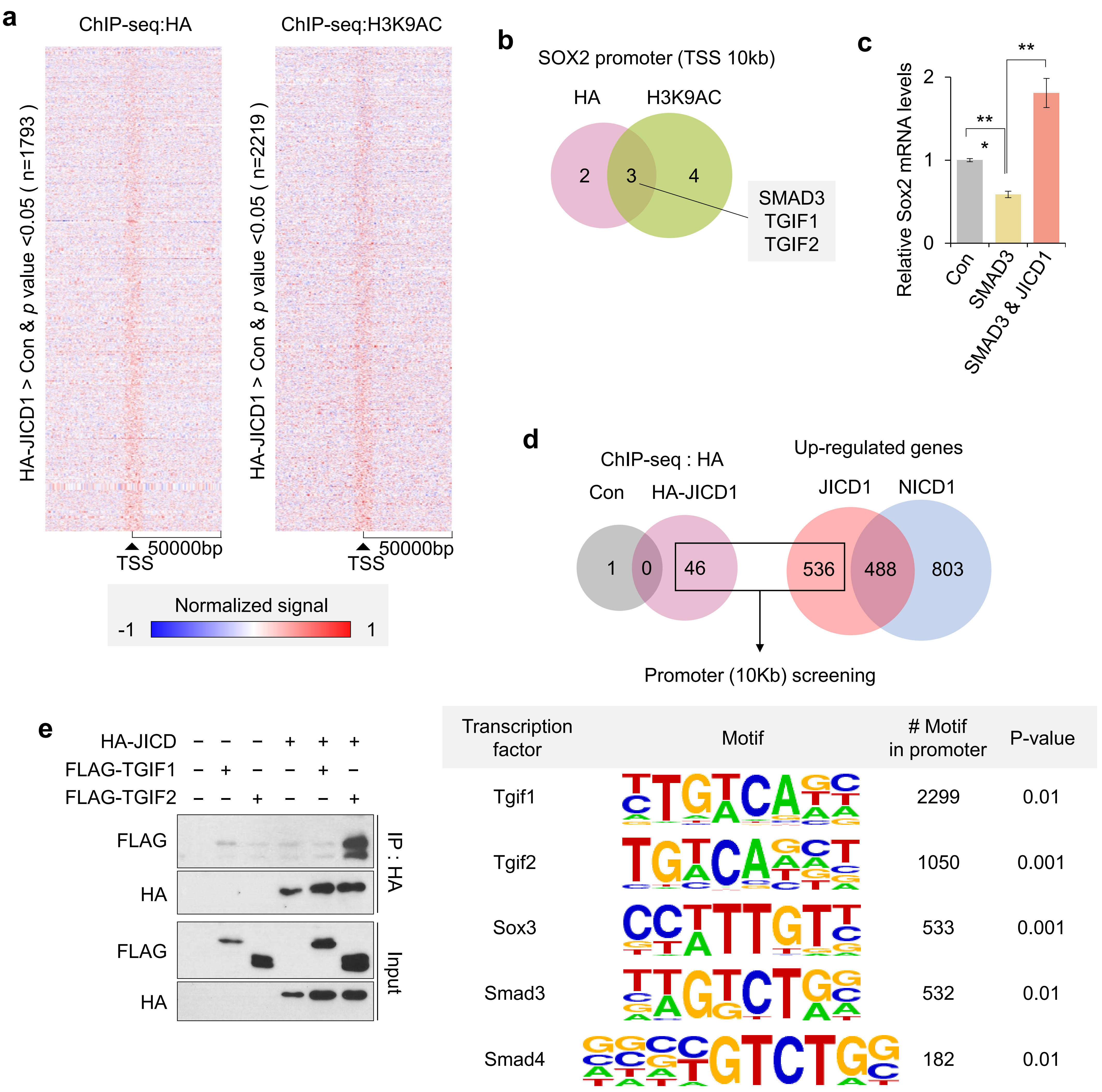


**Supplementary Fig. 9 |** See next page for caption

**Supplementary Fig. 9 | Transcription factors enriched in the promoter of genes upregulated by JICD1 binding. Related to Figures 5 and 6. a**, Heatmaps showing the normalized HA (left) and H3K9Ac (right) ChIP-seq signal for JICD1-overexpressing cells. Numbers indicate the number of genes whose promoter peaks are increased by JICD1. (TSS: Transcription start site). **b**, The transcription factor binding DNA motif commonly detected in ChIP-seq analysis examined using HA and H3K9Ac antibodies at the Sox2 promoter (within -10kb of TSS) in HA-JICD1-overexpressing Ink4a/Arf$^{-/-}$ astrocytes. **c**, Expression of Sox2 mRNA in control, FLAG-SMAD3-, both FLAG-SMAD3-, and HA-JICD1-overexpressing Ink4a/Arf$^{-/-}$ astrocytes. *$p < 0.05$, **$p < 0.01$. **d**, Screening the transcription factor associated with JICD1 in the promoter (within -10kb of TSS) of JICD1-upregulated DEGs. **e**, Co-IP analysis between FLAG-TGIF1/TGIF2 and HA-JICD1 when washed under mild condition.

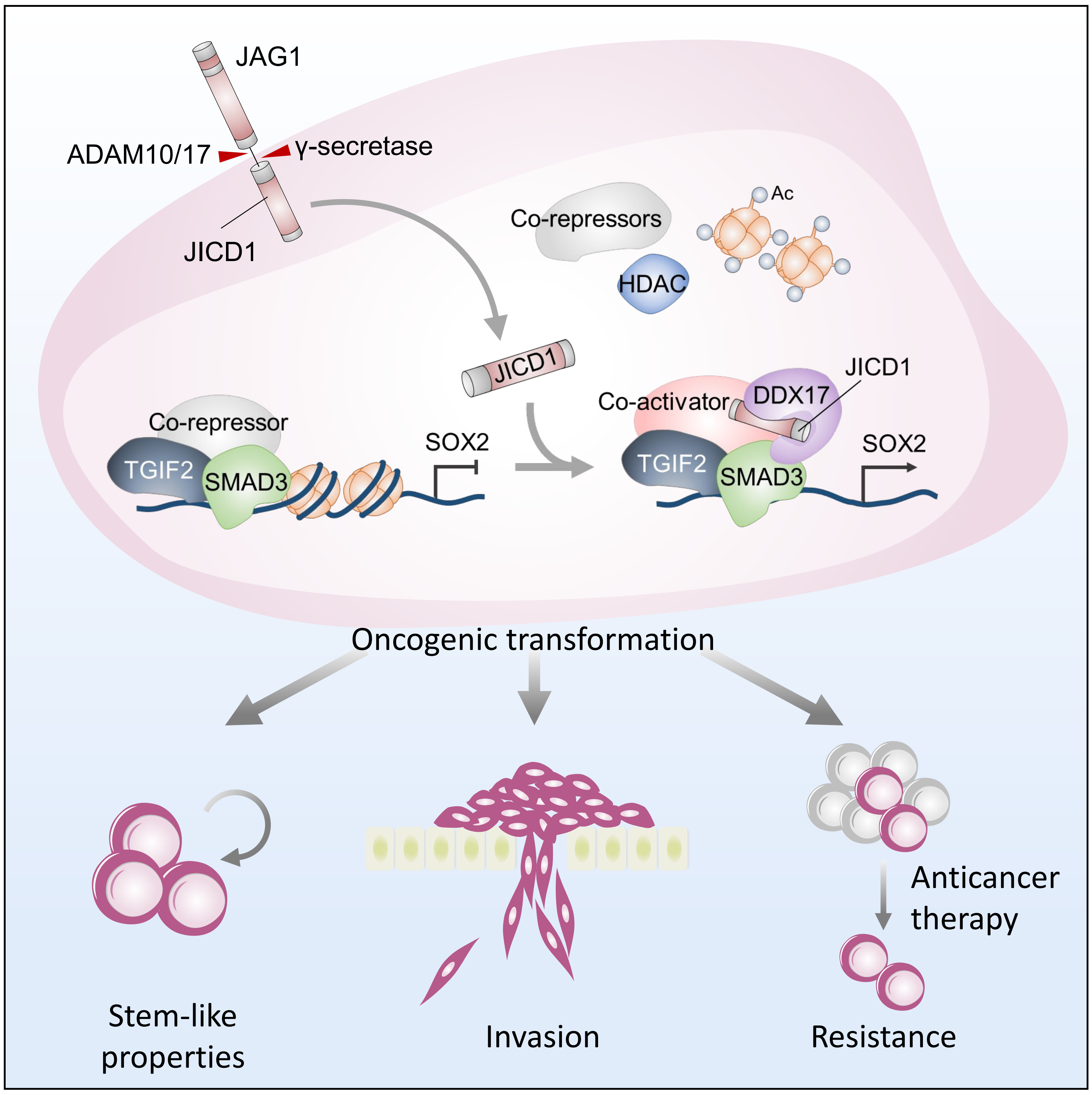


**Supplementary Fig. 10 | Schematic diagram showing the proposed mechanism.** JAG1 intracellular domain generated from JAG1 functions as a transcriptional cofactor in the formation of a transcriptional complex with DDX17, SMAD3, and TGIF2. This complex promotes the expression of SOX2, which plays a crucial role in tumorigenesis by acquiring cancer stem cell properties.
