## Supplementary figures and images for "JAG1 intracellular domain acts as a transcriptional cofactor that forms an oncogenic transcriptional complex with DDX17/SMAD3/TGIF2"

### Kim_JICD_Figure.pdf

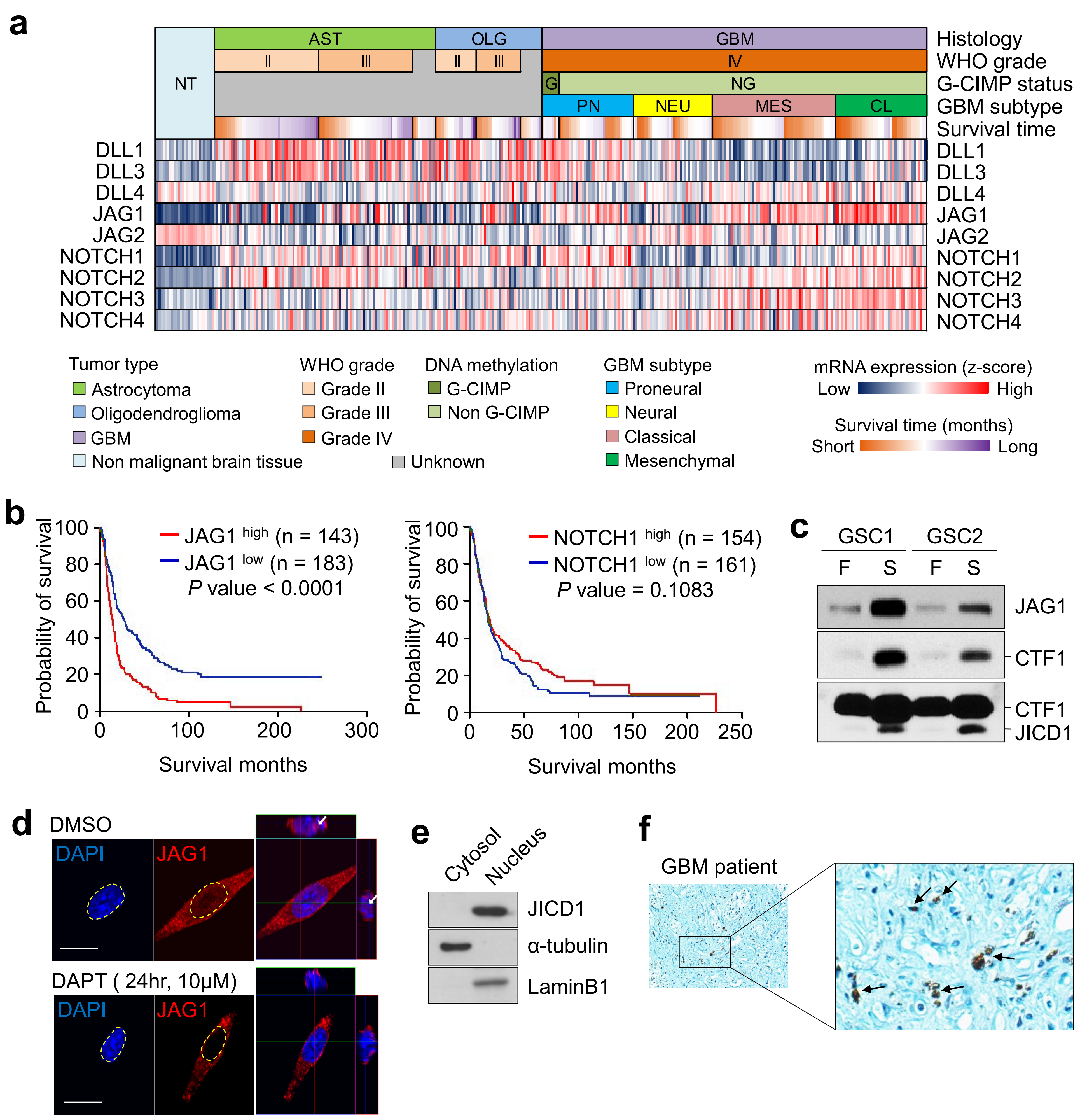

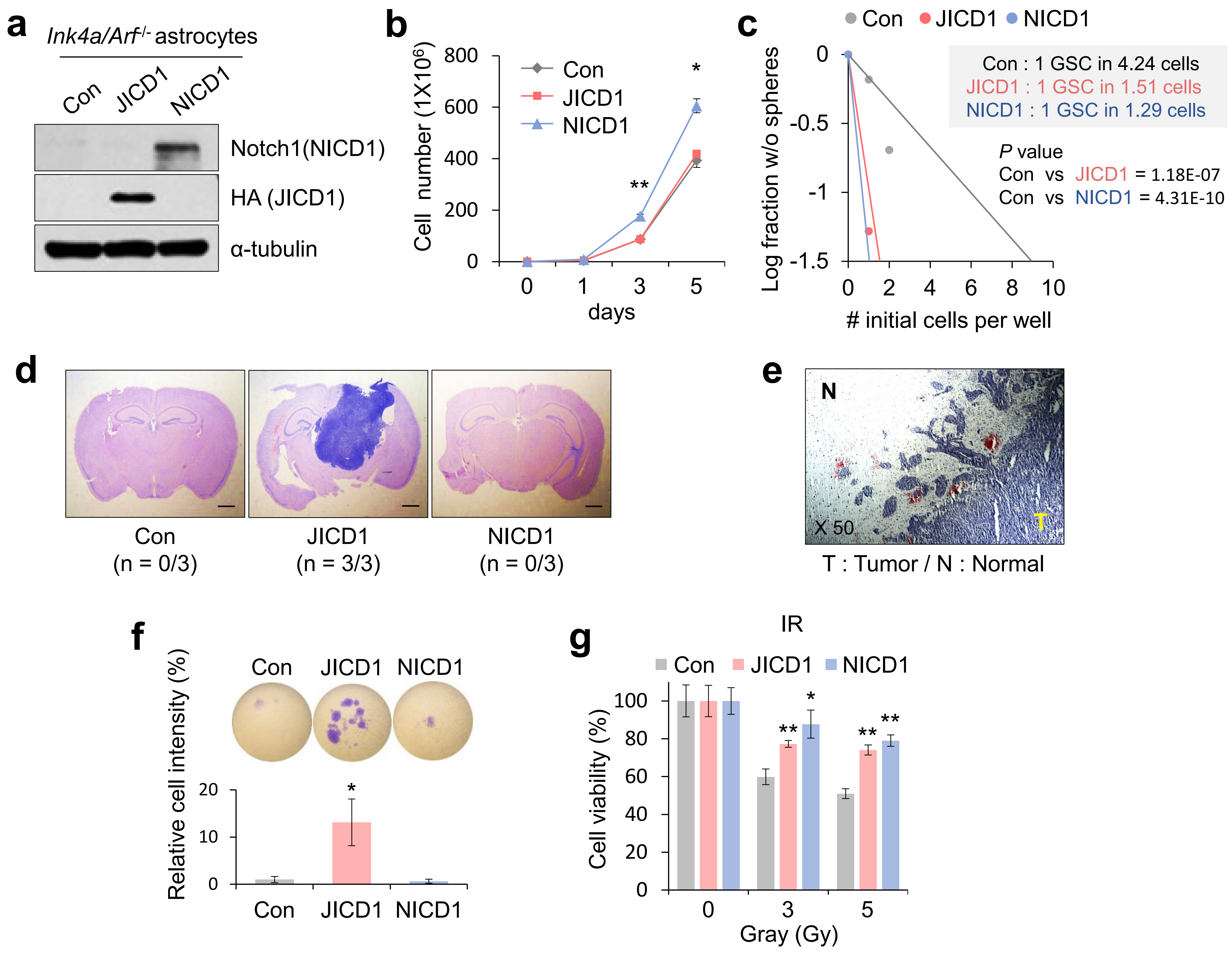

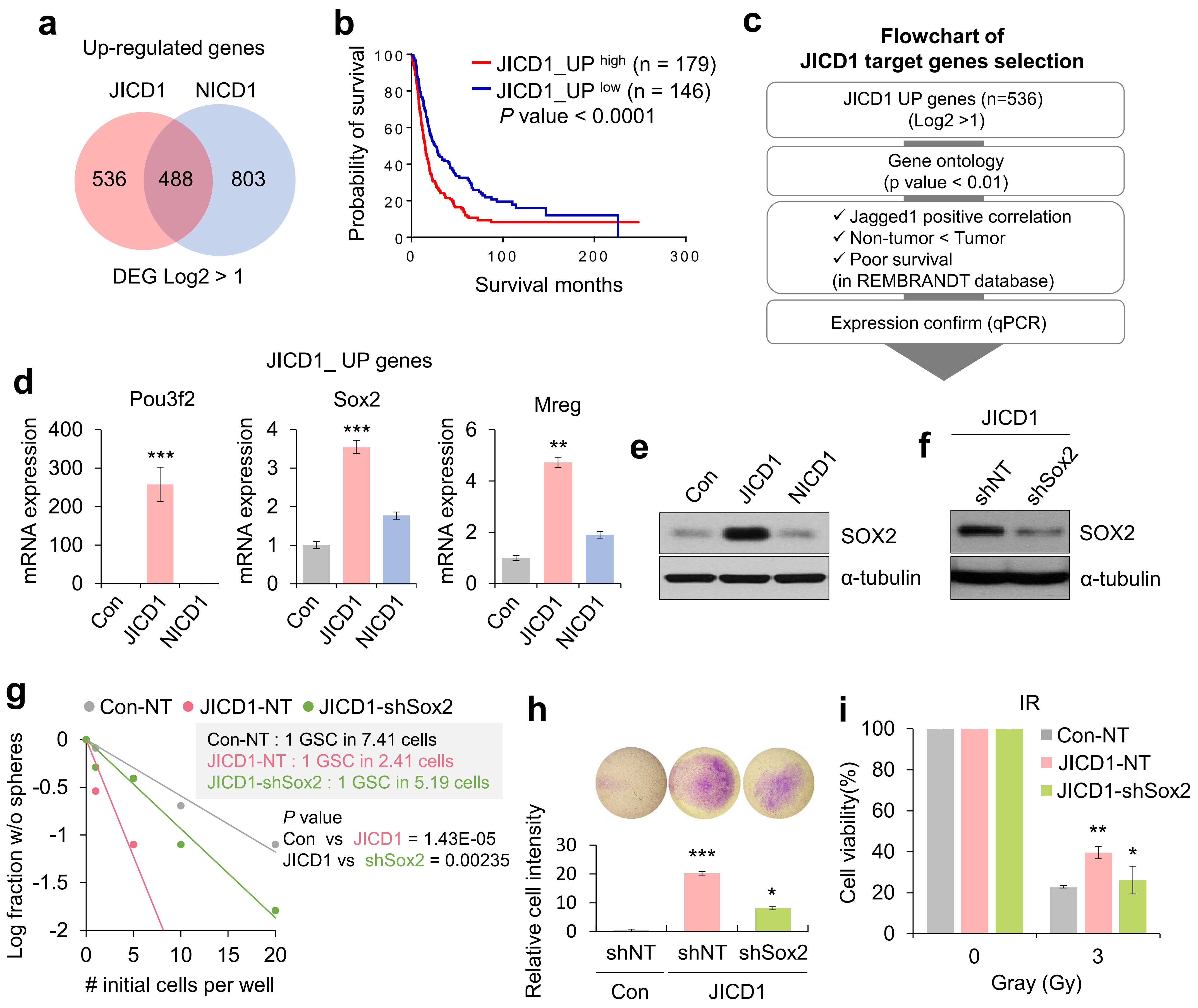

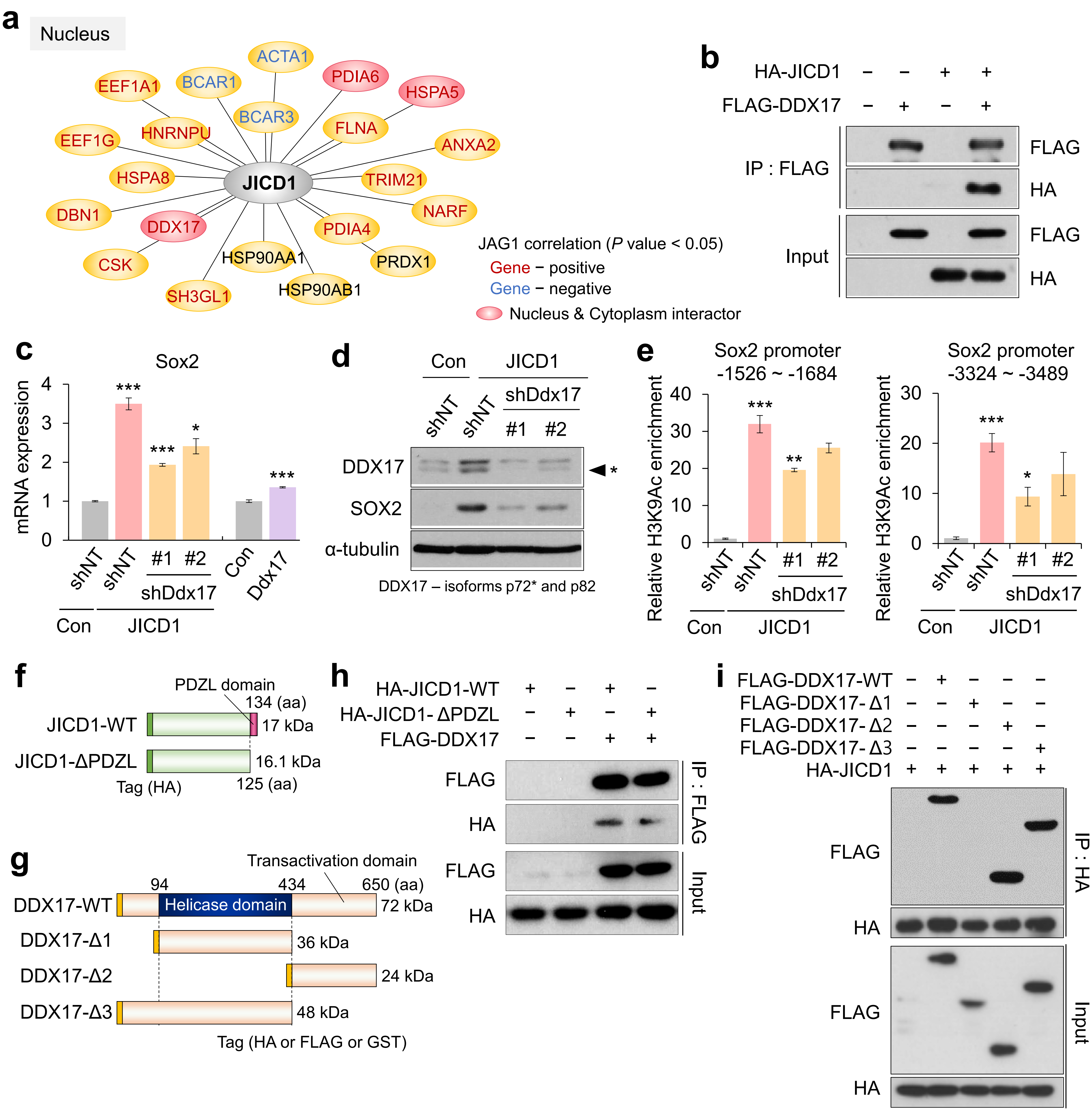

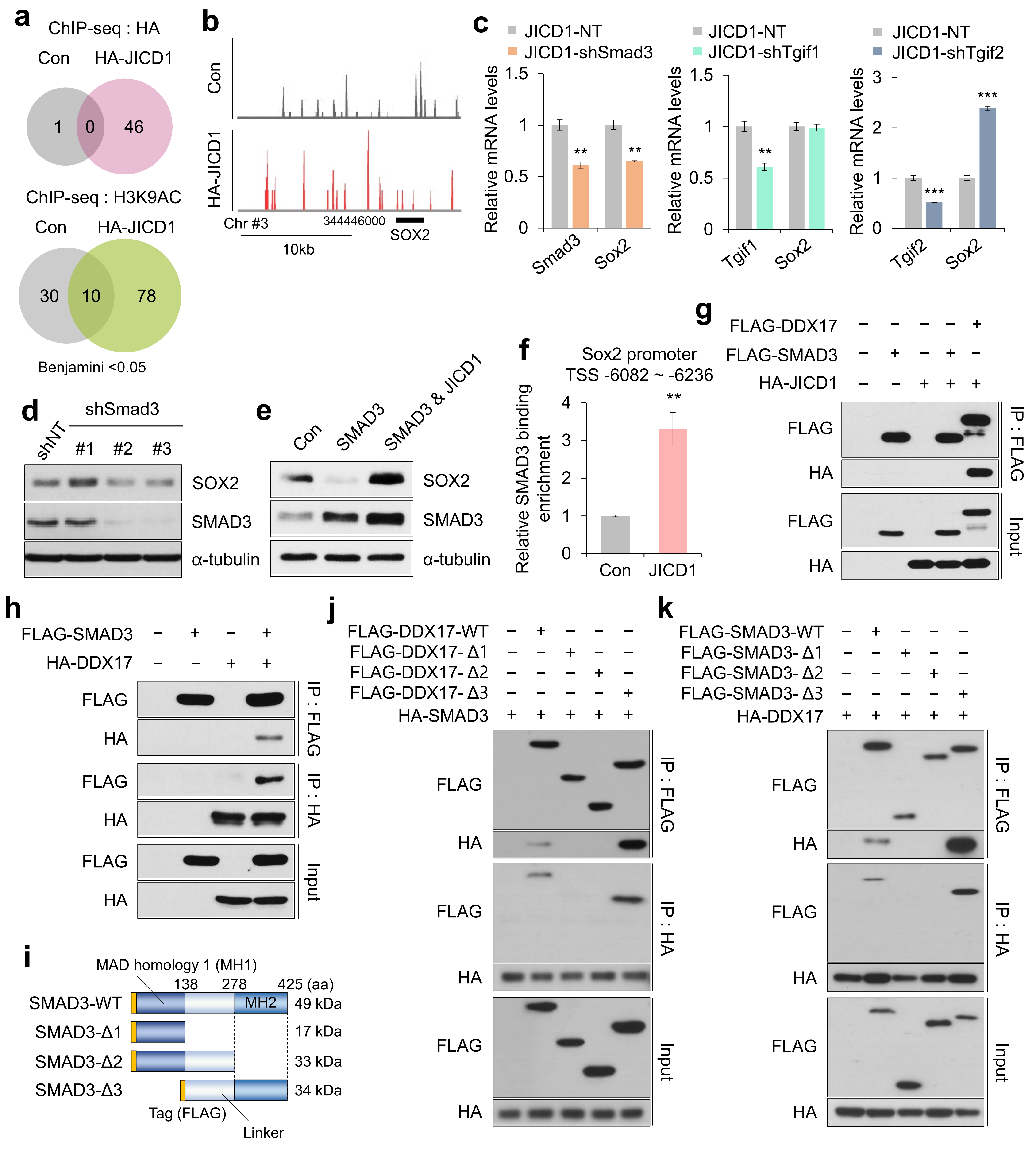

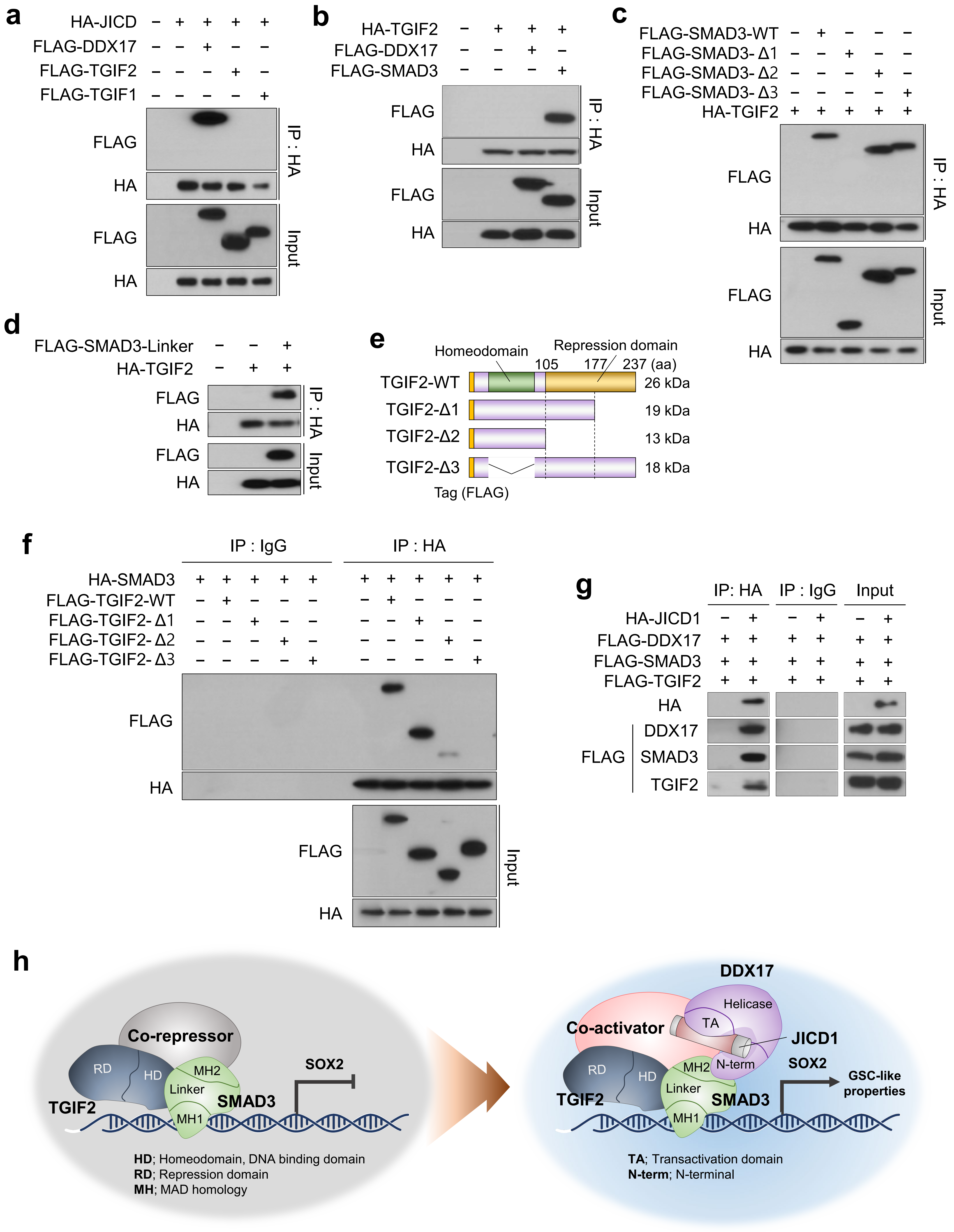
